# RADA is a main branch migration factor in plant mitochondrial recombination and its defect leads to mtDNA instability and cell cycle arrest

**DOI:** 10.1101/856716

**Authors:** Nicolas Chevigny, Frédérique Weber-Lotfi, Cédric Nadiras, Arnaud Fertet, Monique Le Ret, Marc Bichara, Mathieu Erhardt, André Dietrich, Cécile Raynaud, José M. Gualberto

## Abstract

The mitochondria of flowering plants have large and complex genomes whose structure and segregation are modulated by recombination activities. The late steps of mitochondrial recombination are still poorly characterized: while the loss of mitochondrial recombination is not viable, a deficiency in RECG1-dependent branch migration has little impact on plant development, implying the existence of alternative pathways. Here we present *RADA*, an ortholog of bacterial RadA/Sms, which is required for the processing of organellar recombination intermediates. While bacterial RadA is dispensable, RADA-deficient plants are severely impacted in their development and fertility, correlating with increased mtDNA ectopic recombination and replication of recombination-generated subgenomes. The *radA* mutation is epistatic to *recG1*, indicating that RADA drives the main branch migration pathway of plant mitochondria. In contrast, the double mutation *radA recA3* is lethal, underlining the importance of an alternative RECA3-dependent pathway. Although RADA is dually targeted to mitochondria and chloroplasts, we found little to no effects of *radA* on the stability of the plastidial genome. The stunted growth of *radA* mutants could not be correlated with obvious defects in mitochondrial gene expression. Rather, it seems that is partially caused by a retrograde signal that activates nuclear genes repressing cell cycle progression.

## INTRODUCTION

The mitochondrial genomes (mtDNA) of vascular plants are large and complex, mostly consisting of non-coding sequences assembled in a heterogeneous population of linear, circular and branched double-stranded (dsDNA) or single-stranded (ssDNA) DNA molecules (Backert et al., 1997; Bendich, 1996; Manchekar et al., 2006). In most species this collection of subgenomic molecules can be mapped into a single circular chromosome, but multichromosome mitogenomes can also exist (Sloan, 2013). The complexity of the plant mtDNA comes from frequent homologous recombination (HR) events involving repeated sequences. Large repeated sequences (>500 bp) are involved in frequent and reversible HR, while intermediate-size repeats (IRs) (50-500 bp) or microhomologies (<50 bp) can promote infrequent ectopic or illegitimate recombination, respectively (Christensen, 2018; Gualberto et Newton, 2017; Maréchal et Brisson, 2010; Woloszynska et Trojanowski, 2009). Recombination involving IRs or microhomologies contributes to the heteroplasmic state of mtDNA by creating sub-stoichiometric alternative configurations (mitotypes) that co-exist with the main genome (Small et al., 1989). Sub-stoichiometric mtDNA variants can become the predominant genome in the time frame of a single plant generation by a yet unclear process of clonal expansion called sub-stoichiometric shifting (Janska et al., 1998; Small et al., 1989; Vitart et al., 1992). These variants may present altered gene expression, resulting from the displacement of regulatory sequences or from the disruption of gene sequences. Such events may lead to the creation and expression of chimeric ORFs that can be deleterious for mitochondrial function, like in the case of cytoplasmic male sterility (CMS) (Budar et al., 2003; Hanson et Bentolila, 2004). HR is also the main pathway of plant mitochondria for the repair of double strand breaks (DSBs) and the copy-correction of mutations, thus contributing to the very slow evolution of plant mtDNA coding sequences (Christensen, 2013; Mower et al., 2007).

Defect in the factors involved in plant organellar recombination pathways can cause genomic rearrangements, because of increased ectopic recombination resulting from the activation of alternative error-prone repair pathways (Miller-Messmer et al., 2012; Wallet et al., 2015). Many of these factors are derived from prokaryotic homologs inherited from the symbiotic ancestors of mitochondria and chloroplasts (Boesch et al., 2011; Gualberto et Newton, 2017). However, the organellar pathways can significantly depart from thouse of their bacterial counterparts, and involve additional factors with partially redundant functions. As a representative example, plant organellar recombination relies on the abundant RecA-like RECA2 recombinase [(about 450 copies/mitochondrion (Fuchs et al., 2019)], which is targeted to both organelles and whose mutation is lethal at the seedling developmental stage (Miller-Messmer et al., 2012; Shedge et al., 2007). But this process also involves plastidial RECA1 and mitochondrial RECA3. RECA3 could not be detected in Arabidopsis cultured cells (Fuchs et al., 2019), but its function is critical for mtDNA maintenance, since its loss causes mtDNA instability that worsens over generations (Miller-Messmer et al., 2012; Shedge et al., 2007). Double mutants of *RECA3* and other mitochondrial recombination factors such as MSH1 and RECG1 are also much more affected in development than the individual homozygote mutants (Shedge et al., 2007; Wallet et al., 2015). This apparently reflects specialized functions and RECA3-dependant alternative recombination pathways, maybe because *RECA3* is expressed in specific tissues such as pollen and ovules (Miller-Messmer et al., 2012). It was also recently found that tobacco *RECA3* is cell cycle regulated (Trolet et al., 2019).

In bacteria, HR is initiated by the loading of RecA on ssDNA, forming a nucleofilament that then seeks for homologies in the genome by probing multiple heterologous sequences (Forget et Kowalczykowski, 2012; Ragunathan et al., 2012). When a homologous sequence is identified, RecA-mediated ATP hydrolysis stabilizes the invading DNA, forming the synaptic complex also known as displacement loop or D-loop. An important post-synaptic step is branch migration, which involves helicases that extend the homologous region on both sides of the D-loop, allowing the recruitment of the fourth DNA strand to form a Holliday Junction (Cooper et Lovett, 2016; West, 1997; Whitby et al., 1994). Three partially redundant pathways of branch migration have been described in bacteria, involving RuvAB, RecG and RadA, respectively (Beam et al., 2002; Cooper et al., 2015). RadA (also known as “Sms” for “sensitivity to methylmethane sulfonate” and not to be confused with Archea RadA that is unrelated) has long been known as a factor influencing repair by recombination (Beam et al., 2002; Slade et al., 2009), but its biochemical activities were only recently characterized (Cooper et Lovett, 2016; Marie et al., 2017). It is an ATP-dependent ssDNA helicase composed of three functional domains: an N-terminal C4 zinc-finger, a RecA-like ATPase domain and a Lon protease-like domain (Cooper et al., 2015). In contrast to RecG and RuvAB, RadA interacts with RecA and can function in the context of the RecA nucleofilament. Bacterial *radA* or *recG* single mutants are only mildly affected in DNA repair, but the *radA recG* double mutant is severely impaired in its survival under genotoxic conditions (Cooper et al., 2015). Highlighting the crucial role of branch migration in HR, the deficiency in multiple branch migration pathways is more deleterious to the cell than the absence of recombination, because of the accumulation of toxic unprocessed intermediates (Beam et al., 2002; Cooper et al., 2015).

In Arabidopsis mitochondria, RECA-dependent recombination is apparently essential, because *recA2 recA3* double mutants could not be obtained (Miller-Messmer et al., 2012). Plant genomes do not encode any homolog of the RuvAB branch migration complex, but code for an organellar-targeted RecG homolog (RECG1) (Odahara et al., 2015; Wallet et al., 2015). But despite of the absence of a mitochondrial RuvAB pathway, *recG1* mutants are only mildly affected in development. Since mitochondrial HR is apparently essential, a more deleterious effect was expected for the loss of this branch migration activity, suggesting the existence of additional pathways for the maturation of recombination intermediates.

Here we describe a plant homolog to eubacterial RadA/Sms, which could potentially be involved in the late steps of organellar HR pathways. We show that Arabidopsis RADA possesses similar activities as bacterial RadA. However, contrarily to bacteria, plant *radA* mutants are severely affected in their development because of mtDNA instability. The mutation is epistatic to *recG1,* indicating that RADA has a more prominent role in plant mitochondrial recombination than it has in bacteria. Furthermore, we found that mtDNA instability triggered by the deficiency of RADA activates genes involved in the suppression of cell cycle progression, partially explaining the growth defect of *radA* plants.

## RESULTS

### All plant genomes contain a gene coding for RADA, and RADA from Arabidopsis is targeted to both mitochondria and chloroplasts

The Arabidopsis (*Arabidopsis thaliana*) genome was screened for orthologs of RadA and the At5g50340 gene was identified as coding for a protein remarkably similar to bacterial RadA/Sms (45 % similarity), which we named *RADA*. Phylogenetic analysis showed that *RADA* genes are present in all groups of the green lineage, including land plants, green and red algae, as well as in brown algae, diatoms and also in several organisms of the Stramenopile group that are not photosynthetic, such as the water mold *Phytophtora infestans* (Supplemental Figure 1). But no RADA ortholog was found in animals or in yeast. While it is probable that plants inherited *RADA* from the prokaryote ancestor of mitochondria or chloroplasts, the high conservation of the sequences did not allow us to infer whether the ancestor was a proteobacterial or a cyanobacterial RadA. Sequence alignments (Supplemental Figure 2) showed that plant RADA have all the important functional motifs that have been described for bacterial RadA/Sms (Cooper et Lovett, 2016; Marie et al., 2017). In addition, plant RADA sequences have non-conserved N-terminal extensions predicted to be organellar targeting peptides. This was tested by expression of N-terminal fusions to GFP. Fusion proteins were constitutively expressed in transgenic Arabidopsis plants under a strong constitutive promoter and found targeted to both chloroplasts and mitochondria, as shown by co-localization with the autofluorescence of chlorophyll and the red fluorescence of MitoTracker^®^ (Figure 1A–C). The N-terminal part of RADA was sufficient to drive targeting to both organelles (Figure 1B), but the full-length fusion localized in chloroplasts mainly in discrete speckles that could be nucleoids, according to co-localization with PEND:dsRED used as nucleoid marker (Figure 1A and D). A previous report described that, in rice, RADA is targeted to the nucleus (Ishibashi et al., 2006). This was inferred from immunodetection with an antibody raised against the recombinant protein. However, our results did not show any hint that RADA could also be a nuclear protein. Thus, RADA is apparently a dually targeted organellar protein, like RECG1 and several other factors involved in the maintenance of organellar genomes (Gualberto et Newton, 2017).

**Figure 1.**
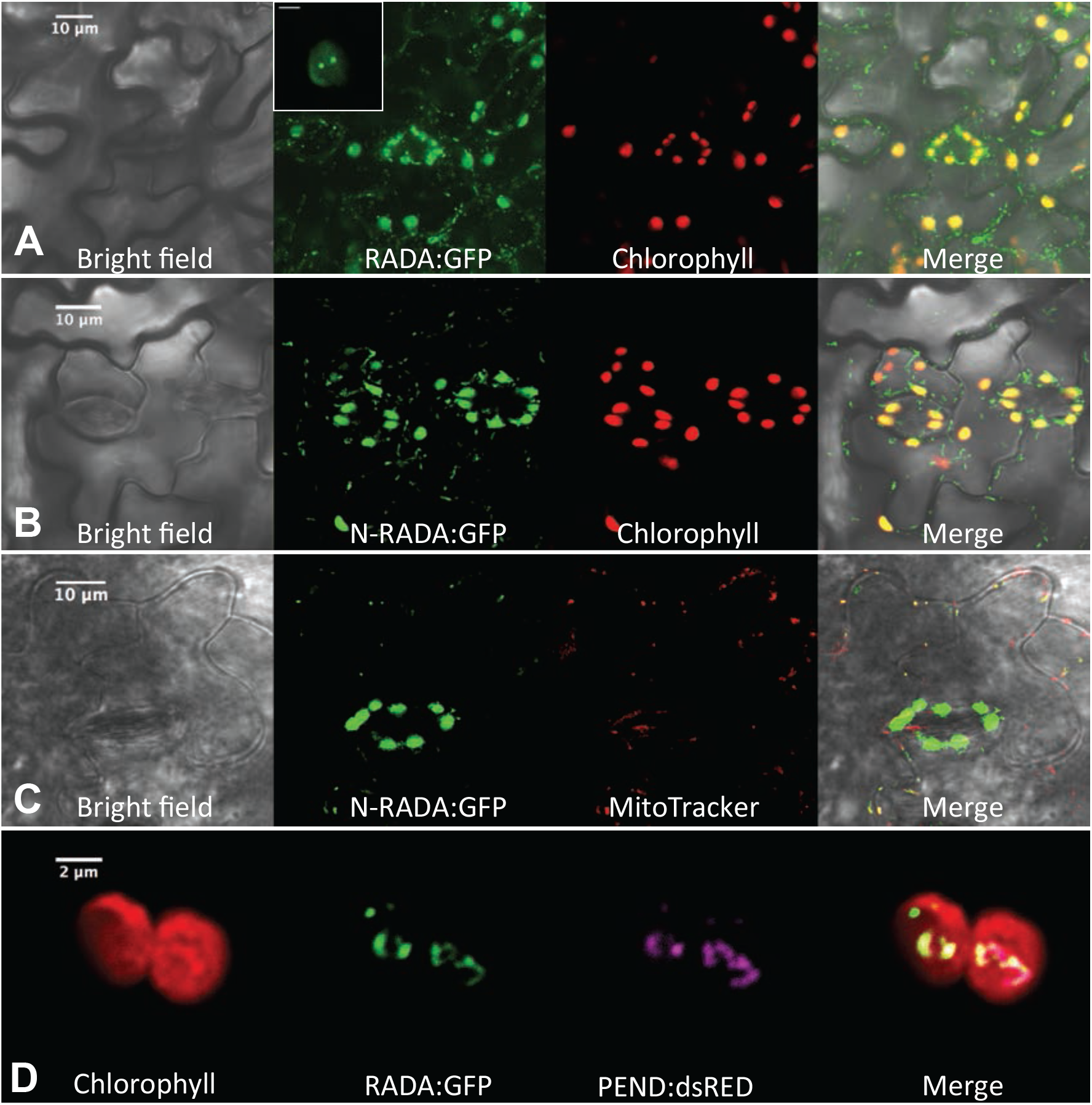
Arabidopsis RADA is targeted to both chloroplasts and mitochondria. (**A** and **B**) Intracellular localization of RADA:GFP fusion proteins constitutively expressed in Arabidopsis transgenic plants. Both the full length protein (RADA:GFP) or just its N-terminal part (N-RADA:GFP) drive localization into chloroplasts (co-localization with autofluorescence of chlorophyll) and into mitochondria (characteristic cytosolic speckles). In chloroplasts RADA:GFP is predominant in dots that could be nucleoids (insert in A). (**C**) Confirmation of co-localization with red fluorescence of MitoTracker. (**D**) Co-localization with plastidial nucleoid marker PEND:dsRED, after biolistic co-transfection in *Nicotiana benthamiana* leaves.

Expression data available at Genevestigator (https://genevestigator.com) indicate that *RADA* is preferentially expressed in hypocotyls and in the shoot apex (Supplemental Figure 3A). In line with these data expression of a promoter:GUS fusion construct was found increased in young shoots, in sepals and in the stigma (Supplemental Figure 3B).

### Structural and functional conservation of plant RADA as compared to bacterial RadA

A three dimensional model of Arabidopsis RADA was built (Figure 2B), based on the known structure of *Streptococcus pneumoniae RadA* (Marie et al., 2017). The RadA structure comprises an N-terminal C4 zinc-finger, required for the interaction with RecA, and two main domains: a RecA-like ATPase domain and a Lon protease-like domain. The RecA-like ATPase domain comprises the Walker A and B motifs, and a RadA-specific KNRFG sequence (Supplemental Figure 2 and Figure 2A). The Walker A and KNRFG motifs are indispensable for the branch migration function of the protein, and are also involved in the DNA-binding and helicase activities of RadA (Cooper et Lovett, 2016; Marie et al., 2017). In bacteria, Walker A and KNRFG mutants are dominant negative, interfering with the function of the wild-type protein. The structural similarity between RadA and RecA suggests functional similarities, but while RecA specialized in the recognition of homologous sequences and strand invasion, RadA rather evolved for driving branch migration. Finally, the C-terminal P-domain of RadA is similar to the Lon protease-like domain of RecA, but it is an inactive domain, since the residues involved in protease activity are not conserved, and no protease activity could be detected (Inoue et al., 2017). Rather, this domain is involved in the binding to DNA and it works as a scaffold for the protein architecture, promoting its oligomerization as hexameric rings (Inoue et al., 2017; Marie et al., 2017).

**Figure 2.**
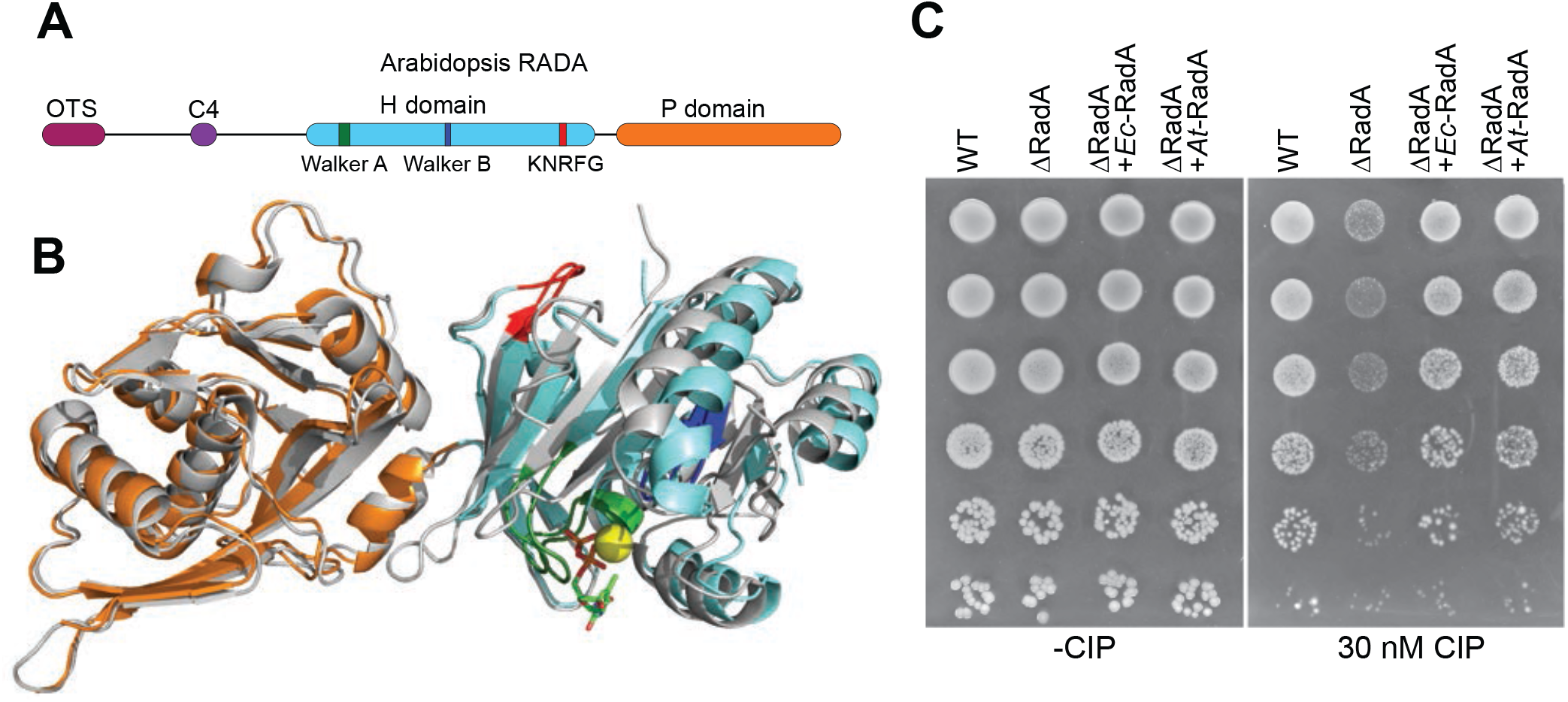
The Arabidopsis RADA is structurally and functionally homologous to bacterial RadA. (**A**) The modular structure of plant RADA is similar to the one from bacteria, with a N-terminal zinc-finger (C4), a helicase domain and a Lon-protease-like domain (H and P domains, respectively). Plant precursor proteins have an N-terminal extension containing an organellar targeting sequence (OTS). (**B**) Model of Arabidopsis RADA superposed on the known structure of *S. pneumoniae* RadA (in gray) (Marie et al., 2017). The color code of relevant domains is as in (A). A bound ADP (stick representation) and the Mg^2+^ ion (yellow ball) are shown. (**C**) Complementation of an *E. coli radA* mutant (ΔRadA) for growth in the presence of genotoxic ciprofloxacin (CIP). Arabidopsis *RADA* (*At*-RADA) complements the mutation as efficiently as the bacterial protein (*Ec*-RadA) cloned in the same expression vector.

We have tested the ability of plant RADA to complement *Escherichia coli* RadA in the repair of genotoxic stress-induced DNA lesions. A *radA785(del)::kan* strain was used for complementation assays, and Arabidopsis RADA and *E. coli* RadA were expressed from the low-copy number plasmid pACYC. As described by others, we found that the *radA* strain was little affected by genotoxic treatments as compared to WT (Beam et al., 2002; Cooper et al., 2015). The conditions we found best to test complementation were in the presence of ciprofloxacin, an inhibitor of gyrase that induces DNA DSBs. In a spot assay, the growth of radA-deficient cells was much reduced as compared to WT ones. Expression of Arabidopsis RADA could complement the ciprofloxacin-triggered growth defect as well as bacterial RadA cloned in the same expression vector (Figure 2C). Therefore plant RADA can functionally substitute bacterial RadA in the repair of DNA damages induced by ciprofloxacin.

### RADA is an ssDNA-binding protein that stimulates branch migration in strand-exchange reactions

The Arabidopsis cDNA sequence minus the first 48 codons coding for the putative organellar targeting sequence was cloned in bacterial expression vector pET28a, fused to an N-terminal His-tag. A mutant sequence was prepared to express a Walker A-deficient protein (K201A). The mutation of the equivalent position in *E. coli* RadA abolished DNA-dependent ATPase activity and generated a dominant negative *radA* allele (Cooper et al., 2015; Cooper et Lovett, 2016). According to the structure of the protein bound to ADP the mutation should not affect ATP or ADP binding, but ATPase activity is lost (Supplemental Figure 4).

Both WT RADA and K201A could be expressed and purified as soluble proteins (Supplemental Figure 5A). By gel filtration the purified WT protein resolved as two peaks of high molecular weight (Supplemental Figure 5B), indicating different degrees of protein oligomerization. Dynamic light scattering analysis of the smaller size oligomer showed that it was mainly constituted by a monodispersed particle of about 340 kDa, consistent with a hexameric RADA complex (Supplemental Figure 5C). EMSA experiments showed that both peak fractions could bind to an ssDNA oligonucleotide probe and give rise to complexes of same apparent size (Supplemental Figure 5D).

The purified proteins were tested in EMSA experiments with different DNA structures as substrates, including ssDNA, dsDNA, fork-like structures and double-stranded molecules containing 5’ or 3’ ssDNA overhangs. Using the same probe concentrations and increasing concentrations of recombinant protein we found that RADA could bind to all structures containing ssDNA regions. It could also bind dsDNA, but with much less affinity (Figure 3A). In our experiments we found that higher molecular weight complexes could be formed. These could not be well resolved in standard gel migration conditions, but in 4.5 % polyacrylamide gels and at lower voltage we could resolve these complexes formed with ssDNA, and found that they were promoted by the presence of ATP or ADP (Figure 3B). Such higher molecular weight complexes could correspond to the polymerization of RADA on ssDNA, forming nucleofilaments. We did not see significant differences in binding with the K201A mutant protein, which seemed to bind ssDNA with equivalent affinity as the WT protein. The sequence specificity of RADA binding to ssDNA was tested in competition experiments. In these experiments poly-dT, poly-dC and poly-dG could compete binding as efficiently as the homologous probe that was used, while poly-dA was less efficient as a competitor (Figure 3C). Thus, Arabidopsis RADA binds preferentially to ssDNA, with little sequence specificity.

**Figure 3.**
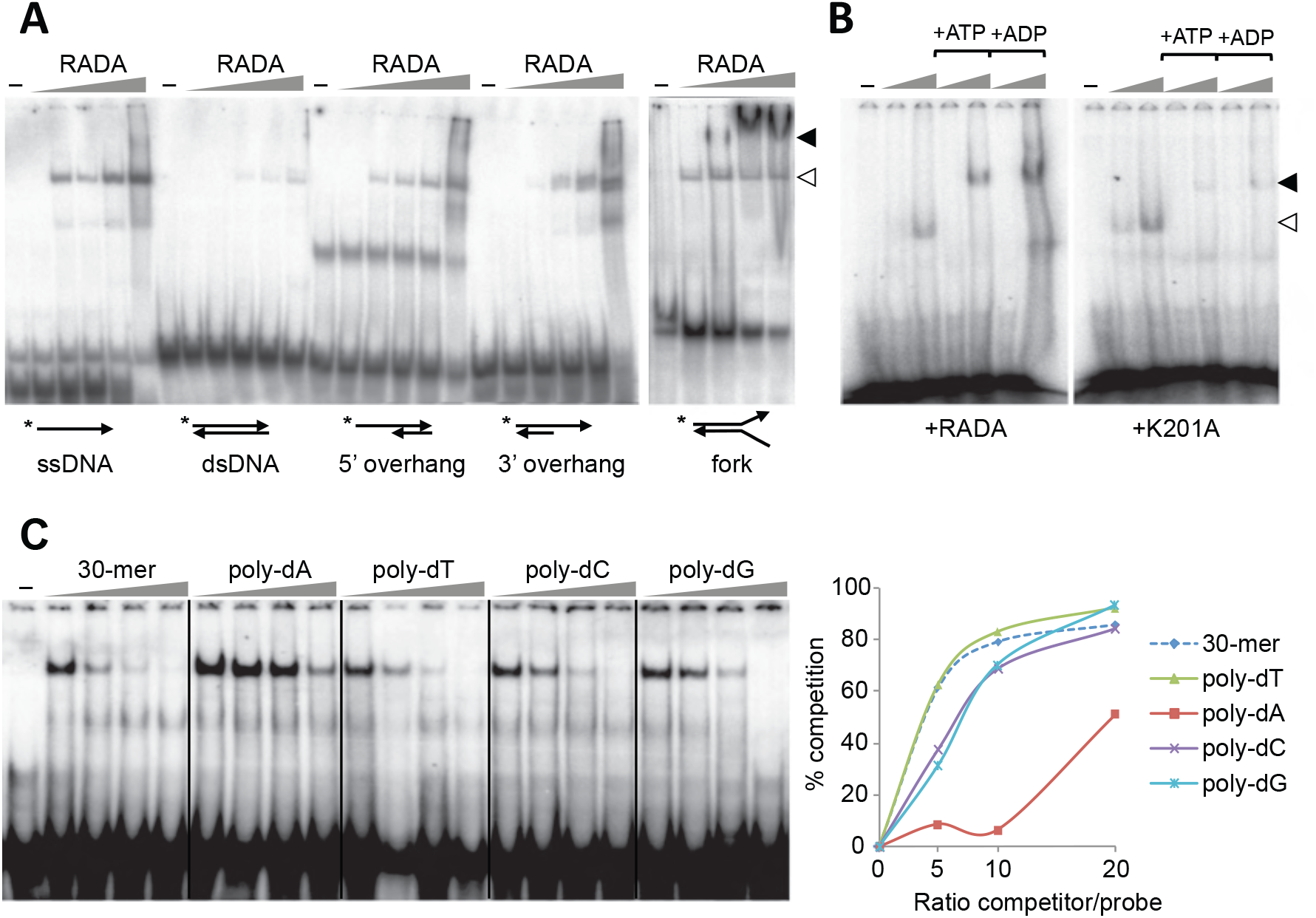
RADA preferentially binds ssDNA. (**A**) EMSA experiments showing that RADA binds any ssDNA-containing DNA structure with higher affinity than dsDNA. Lower and higher molecular weigh complexes are indicated by white and black arrowheads respectively. A predominant band seen with the 5’-overhang probe is artefactual, already present in the absence of protein. (**B**) Analysis on low concentration gel (4.5 % as compared to 8 % in A) of the formation of a high-molecular weight RADA filament on ssDNA, which is promoted by ATP or ADP (1 mM). The K201A mutant protein binds ssDNA with an affinity comparable to the one of the WT protein. Increasing concentrations of RADA or of K201A used in (A) and (B) are indicated by the grey triangles. (**C**) Competition experiments. A 30-mer ssDNA oligonucleotide (7x[AGTC]AG) was used as probe in EMSA experiments with recombinant RADA, and sequence specificity was tested by competition with increasing concentrations of the cold homologous oligonucleotide or with 30-oligomers (poly-dA, pol-dT, poly-dC and poly-dG). Quantification of the results is shown on the right. Only poly-dA showed reduced competition for RADA binding to ssDNA.

The branch migration activity of plant RADA was also tested, using an *in vitro* strand-exchange reaction promoted by bacterial RecA. RecA, in the presence of ATP and bacterial single-strand binding protein SSB, can initiate the invasion of dsDNA by homologous ssDNA and promote branch migration till the final heteroduplex product is completed. In the presence of plant RADA the branched intermediates were resolved faster, leading to an earlier appearance of the final nicked double-stranded circular product (Figure 4A). The faster resolution of recombination intermediates was reproducibly observed in six independent experiments (Figure 4B).

**Figure 4.**
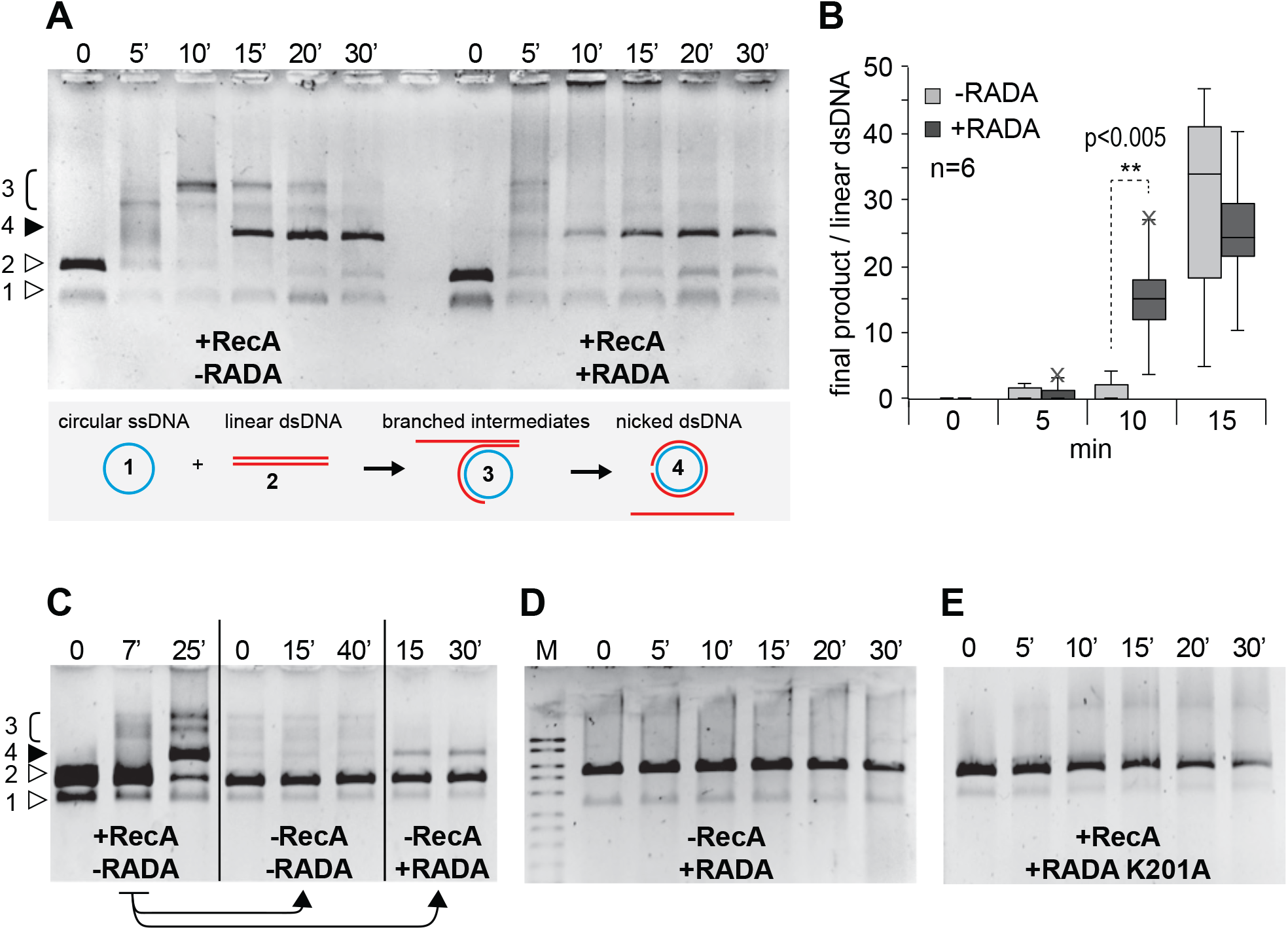
Branch-migration activities of RADA. (**A**) In an *in vitro* strand-invasion reaction recombinant RADA accelerates branch-migration of DNA heteroduplexes initiated by RecA. An explanation of the different substrates and products is shown. (**B**) Ratio of final product as compared to the initial linear dsDNA substrate in 6 independent experiments, showing that in the presence of RADA there is faster resolution of branched intermediates. (**C**) Experiment showing that RADA alone can finalize branch-migration initiated by RecA: a reaction at t=7 min was arrested by deproteination (left panel) and the DNA purified. Without further addition of RecA or RADA proteins there is no spontaneous progression of the reaction (middle panel), but added RADA can alone resolve the recombination intermediates into the final product (right panel). (**D**) RADA alone cannot initiate strand invasion. (**E**) Mutation of the ATPase Walker domain of RADA (K201A) inhibits the reaction.

To test whether RADA-promoted branch migration requires interaction with RecA, assays were arrested by freezing at a time point (7 min) when there was already accumulation of branched intermediates, but no visible final heteroduplex product (Figure 4C left panel). The reaction mix was deproteinated and the purified nucleic acids added to new reaction mixes, in the presence or absence of RADA. In the absence of both RecA and RADA the branched intermediates could not spontaneously evolve and remained stable (Figure 4C middle panel), but in the presence of RADA they were converted to the final product, showing that plant RADA alone can promote branch migration (Figure 4C right panel). This departs from what was described for bacterial RadA, whose function seems to depend from its interaction with RecA. However, RADA is not a recombinase redundant with RecA, because RADA alone in the absence of RecA is not able to initiate the strand-invasion reaction (Figure 4D). This result diverges from a previous report that claimed that plant RADA is able to promote invasion of dsDNA by homologous ssDNA (Ishibashi et al., 2006). Finally, in the same experimental conditions the K201A mutant protein was unable to promote branch migration, and rather completely inhibited the reaction (Figure 4E). That could be because the incapacity of the mutant protein to migrate along the heteroduplexes blocks the activity of RecA. The result is consistent with the dominant negative effect of the equivalent mutation in *E. coli* RadA (Cooper et al., 2015).

### Arabidopsis plants deficient in RADA are severely affected in their development and fertility

We could retrieve several potential *radA* loss-of-function (KO) lines from available Arabidopsis T-DNA insertion collections, from which two (*radA-1* and *radA-2*) could be validated, both in the Col-0 background, with T-DNA insertions in exons 8 and 5 respectively (Figure 5A). Homozygous plants from both mutant lines displayed severe phenotypes of retarded growth of both leaves and roots, and of distorted leaves showing chlorotic sectors (Figure 5B-D). The phenotype was fully penetrant, with all homozygous mutants displaying the phenotype. Transmission electron microscopy (TEM) showed that *radA* mesophyll cells were almost indistinguishable from WT, with chloroplasts that were morphologically normal. On the other hand, mitochondria looked enlarged in size, and less electron dense as compared to those from WT leaves (Supplemental Figure 6). This suggested that it is the mitochondrion that is predominantly affected in *radA*. The severe dwarf phenotype could be partially relieved by growing *radA* plants under a short day photoperiod (8h light/16h dark). In such conditions plants revealed extended juvenility, with dramatic elongation of lifespan and development of aerial rosettes (Figure 5E). A similar phenotype of perennial growth under short days was described for Arabidopsis *msh1* mutants, deficient in the homolog of bacterial mismatch repair protein MutS and affected in mtDNA stability (Xu et al., 2012).

**Figure 5.**
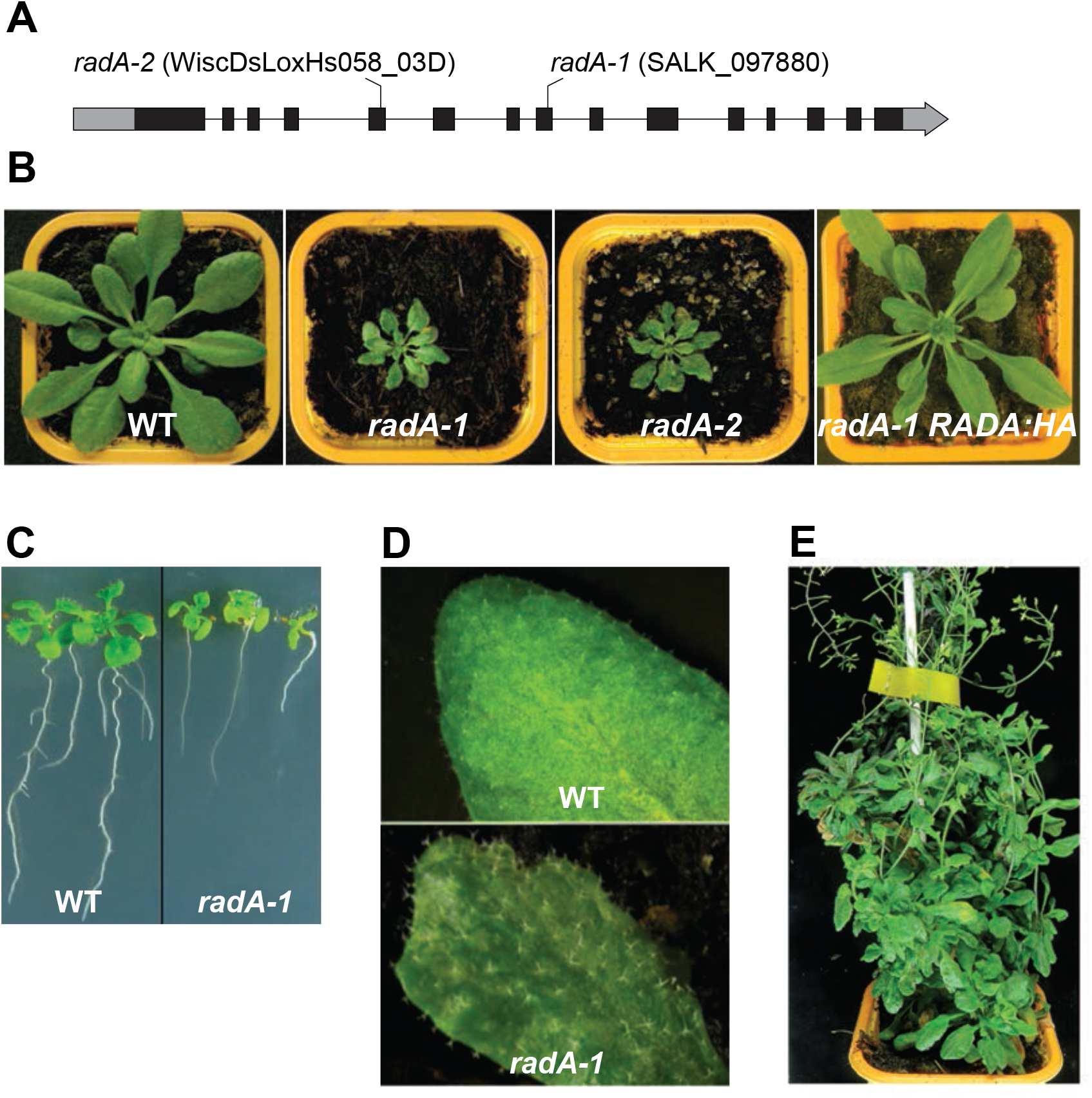
Arabidopsis *radA* mutants and phenotypes. (**A**) Schematic representation of the Arabidopsis *RADA* gene. Coding sequences are in black and 5’- and 3’-UTRs are in gray. The position of the T-DNA insertions in *radA* mutant lines is shown. (**B**) *radA* plants show severe growth retardation, with distorted leaves presenting chlorotic sectors. These phenotypes can be complemented by expression of HA-tagged RADA (RADA:HA) under control of the endogenous *pRADA* promoter. (**C**) root shortening of *radA* plants. (**D**) Detail of leaf phenotype. (**E**) 4-month old *radA-1* plant grown under short days (8h light), showing perennial vegetative growth with development of aerial rosettes.

To fully confirm that these phenotypes were because of a deficiency in RADA, *radA-1* plants were complemented with the WT *RADA* gene expressed under its own promoter. Heterozygote *radA-1* plants were transformed and homozygous plants that also contained WT *RADA* as a transgene were segregated in the T2 generation. These were phenotypically normal (Figure 5B), confirming the complementation and the linkage of the growth defects to the *radA* allele.

Arabidopsis *radA* produces flower stems with very small siliques that mainly contain aborted seeds. The few seeds produced are heterogeneous in shape and few are able to germinate (Figure 6A and 6B). Pollen stained positive by Alexander staining, suggesting that it was viable. That was confirmed by the successful transmission of the *radA* allele in crosses using *radA* pollen. But pollen production was much reduced compared to WT, and many pollen grains were of aberrant size and shape (Figure 6C). Regarding female organs, the stigma displayed elongated and fully differentiated papillae, but with no attached pollen grains (Figure 6D). That could be because papillae cells are modified and unable to bind pollen, or because the stigma develops and is receptive before the pollen matures. To test whether *radA* female gametes are viable emasculated flowers from WT or *radA* were pollinated with WT pollen and ovules were observed before and after pollination. The mature unfertilized *radA* ovules looked morphologically normal and alike WT ovules (Figure 6E). Three days after pollination virtually all WT ovules were fertilized and showed a developing embryo, but only 1/6th of the *radA* ovules had developing embryos (Figure 6E). No elongation of the pollinated pistils was observed even at seven days after pollination (Figure 6F) showing that normal pollen could not fertilize the apparently normal *radA* ovules, potentially because of a deficiency in pollen germination in the stigma of *radA* plants. It is tempting to speculate that it is because of enhanced expression of RADA in the stigmata, as suggested by the promoter:GUS results. Thus, the partial sterility of *radA* would be due to both male and female defects.

**Figure 6.**
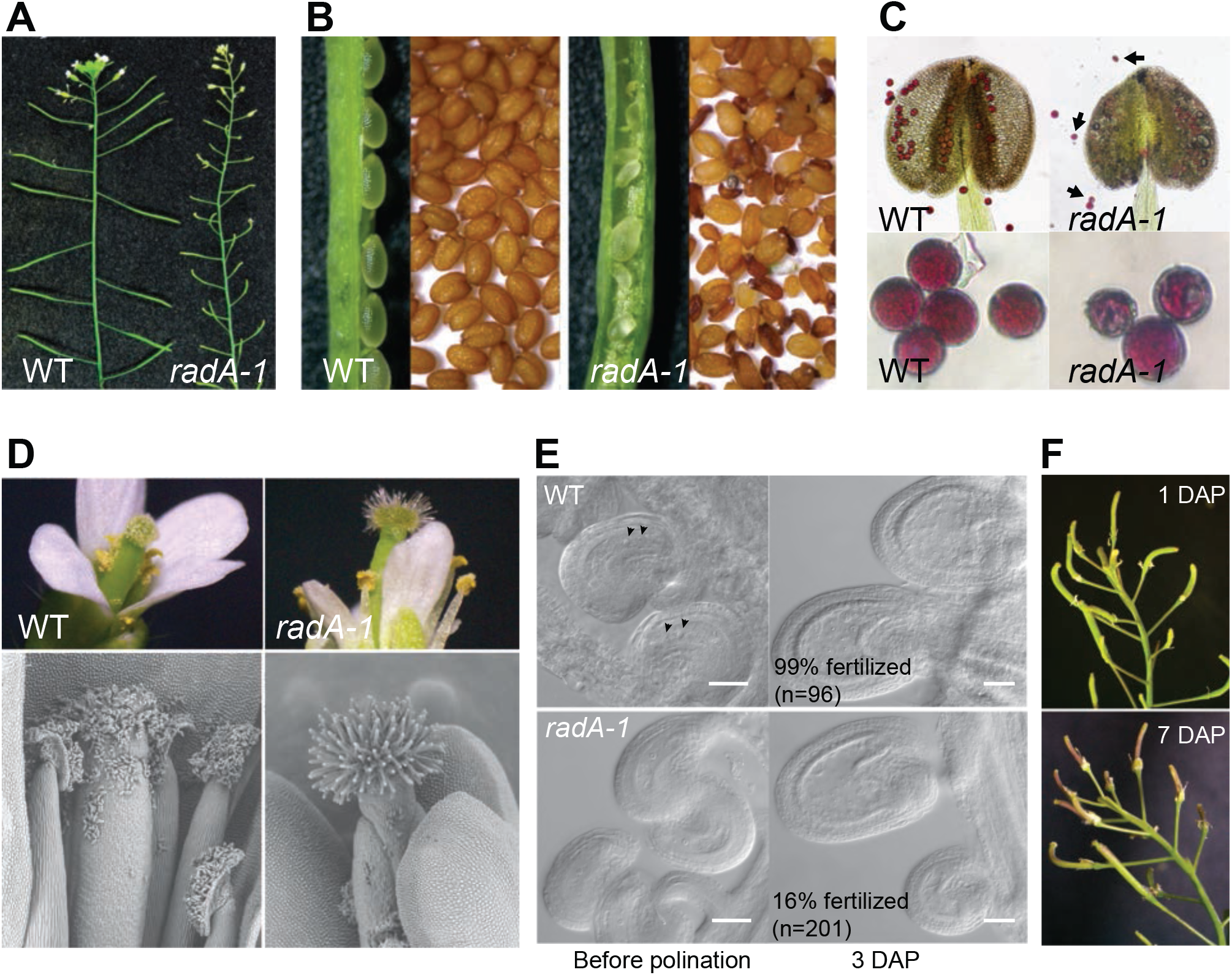
Reduced fertility of *radA* plants. (**A**) Comparison of WT and *radA* flower stems showing very small *radA* siliques. (**B**) *radA* siliques mostly contain aborted seeds, and most of the few seeds produced are non-viable. (**C**) Alexander staining of pollen in *radA* anthers as compared to WT, showing little pollen production and an abundance of small and aberrant pollen grains (indicated by arrows). (**D**) Visible and SEM Images showing that in *radA* no pollen binds to the papillae of the stigma. (**E**) Differential interference contrast images of ovules in crosses between *radA-1* flowers and WT pollen. Black arrowheads indicate central cell and egg cell nuclei in unfertilized ovules. White arrowheads indicate developing embryos in fertilized ovules, at the 2 up to 8 cells globular stage, three days after pollination (DAP). Only 16 % of *radA* ovules could be fertilized. Scale bar is 50 μm. (**F**) At seven DAP the pollinated pistils did not develop further.

### *radA* mutants are affected in the stability of the mitochondrial genome

Several Arabidopsis mutants affected in recombination functions (ex: *msh1, osb1, recA2*, *recA3, recG1*) show, in normal growth conditions, increased ectopic recombination of the mtDNA across intermediate-size repeats (IRs), and the severity of the molecular phenotype normally correlates with the severity of the developmental phenotypes (Arrieta-Montiel et al., 2009; Gualberto et Newton, 2017; Zaegel et al., 2006). The *radA* mutants were thus also tested for such molecular phenotype. Plants from both *radA-1* and *radA-2* lines of the second homozygous generation were grown *in vitro* and four were selected according to the severity of growth defect phenotype (Figure 7A). The relative copy number of the different mtDNA regions in these plants was quantified by qPCR, using a set of primer pairs spaced about 5 kb apart across the genome, as described (Wallet et al., 2015). Dramatic changes in the stoichiometry of mtDNA sequences were observed in all plants (Figure 7B). These affected similar regions of the genome in the individual plants, but the amplitude of the changes was higher in the severely affected plants (*radA-1 #1* and *radA-2#1*) than in the mildly affected ones (*radA-1#2* and *radA-2#2*). An increase in copy number of large genomic regions could be observed, that could be as high as seven fold. Several of the observed events of stoichiometry variation corresponded to regions flanked by pairs of directly oriented repeats, including the pair of repeats A (556 bp), F (350 bp), L (249 bp), and EE (127 bp) (Figure 7B). This suggested that the process at play was the same as that described for repeat EE in the *RECG1* KO mutant, *i.e.* the looping out of a circular subgenome by recombination across directly oriented IRs, followed by its autonomous replication (Wallet et al., 2015). We tested by qPCR the accumulation of crossover products for repeats F, L and EE, as previously described (Miller-Messmer et al., 2012; Wallet et al., 2015). As expected, in all plants a significant increase in crossover products *versus* WT levels was observed for all analyzed repeats (Figure 7C), with a significantly higher accumulation in the more affected plants than in the mildly affected ones. Recombination resulted in the asymmetrical accumulation of mainly one of the two crossover products, with the remarkable exception of recombination involving the pair of repeats L, which resulted in the accumulation of only one of the reciprocal crossover products in mildly affected plants, while both products accumulated in the severely affected plants. As described before, asymmetric recombination could be because of repair by the break-induced replication (BIR) pathway (Christensen, 2018; Gualberto et Newton, 2017). Big differences in the relative accumulation of recombination products were seen, depending on the pair of repeats analyzed. But these values are misleading, because compared to the basal levels that exist in WT plants. Thus, a 30 fold increase in recombination product L-1/2 might be equivalent to a 1000 fold increase in product EE-2/1, because the former is already quite abundant in WT Col-0 and easily detected by hybridization, while the latter is virtually absent in WT (Wallet et al., 2015; Zaegel et al., 2006). Analysis of recombination involving repeats A was not possible, because the size of the region to be amplified (larger than 600 bp) is not compatible with qPCR.

**Figure 7.**
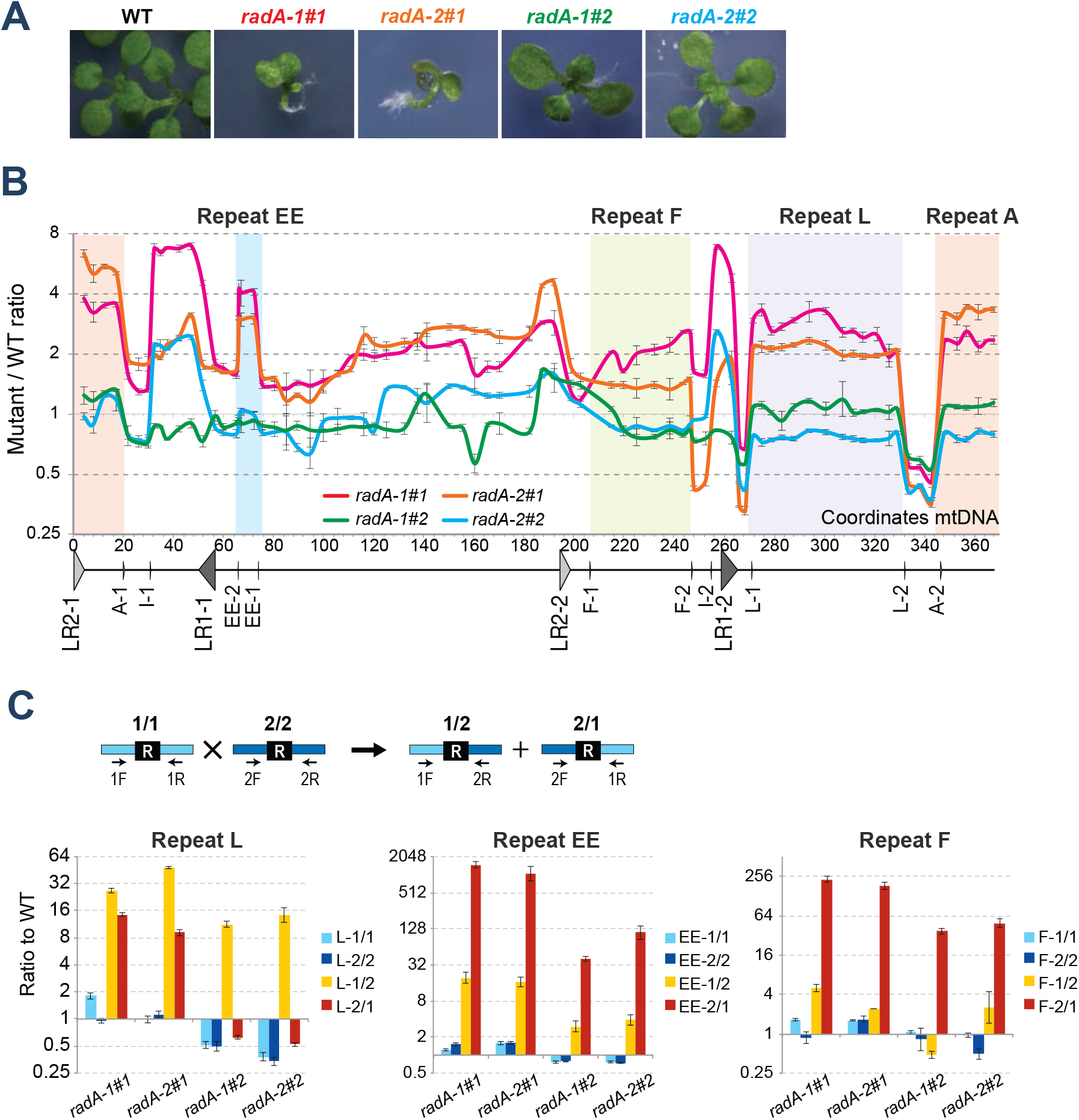
Changes in mtDNA sequences stoichiometry in *radA* mutants because of increased ectopic recombination across repeats. (**A**) Picture of severely affected (*radA-1*#1 and *radA-2*#1) and mildly affected (*radA-1*#2 and *radA-2*#2) seedlings. (**B**) Scanning of their mtDNA for changes in relative copy numbers of the different mtDNA regions. Sequences spaced 5-10 kb apart on the mtDNA were quantified by qPCR. Coordinates are those of the Col-0 mtDNA sequence. The position of the mtDNA large repeats LR1 and LR2 and of relevant intermediate size repeats are shown below the graphic. Regions with changed stoichiometry flanked by repeat pairs are shadowed. (**C**) Accumulation of crossover products from mtDNA repeats L, F and EE in *radA* seedlings as compared to WT. Results are in a log2 scale. The scheme above shows the qPCR relative quantification of parental sequences 1/1 and 2/2 and of the corresponding crossover products 1/2 and 2/1. Results are the mean of three technical replicates, and error bars correspond to *SD* values.

Because RADA is also targeted to chloroplasts the plastidial DNA (cpDNA) of *radA* plants was likewise scanned for changes in sequence stoichiometry by qPCR. No changes were detected between mutant and WT, apart from a slight general increase in cpDNA copy number in several individual plants, but below two-fold and probably not significant. We confirmed this result by Illumina DNAseq of four individual *radA-1* plants. The normalized coverage of the cpDNA did not reveal any changes in sequences stoichiometry as compared to WT plants (Figure 8A). No major proportion of indels was detected neither. We also checked whether *radA* plants could have increased rearrangements of the cpDNA, as described for Arabidopsis *why1why3* and *why1why3reca1* Arabidopsis mutants, and for maize *cptK1* (Duan et al., 2020; Le Ret et al., 2018; Zampini et al., 2015). Analysis by PCR using divergent primers has been used before to detect increased rearrangements of the cpDNA (Duan et al., 2020; Marechal et al., 2009). But in our hands, after testing six pairs of primers described in the literature we just obtained either negative or irreproducible results. We have therefore developed a pipeline for identification of Illumina reads corresponding to cpDNA rearrangements (Supplemental Figure 7). The *radA* sequence libraries contained a slightly higher proportion of rearrangements as compared to WT, but the factor was below two-fold (Figure 8B), *i.e.* too low to be considered as responsible for the severe growth defect of *radA* plants. Thus, in contrast to the major effects observed on mtDNA maintenance, the loss of RADA apparently does not significantly affect the stability of the cpDNA in Arabidopsis.

**Figure 8.**
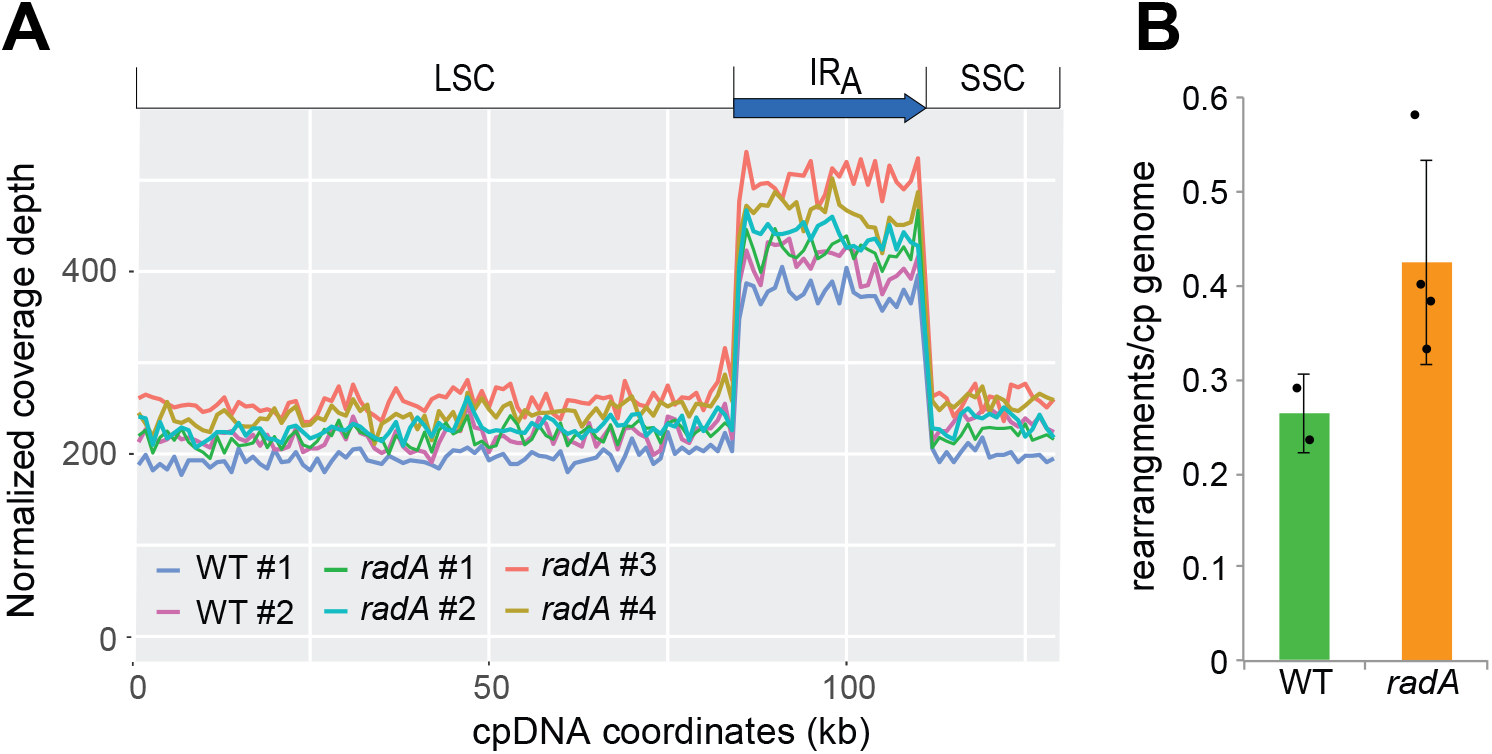
No significant effect of the *radA* mutation on the stability of the plastidial genome. The DNA from a leaf of four independent 4-week-old *radA-1* plants and of two WT plants was extracted and sequenced on a MySeq Illumina system (2×150 bp paired-end). The reads corresponding to the cpDNA were extracted and investigated for possible changes in cpDNA stability. (**A**) Normalized cpDNA coverage, showing comparable copy number of the cpDNA in WT and in *radA*, and no apparent changes in sequences stoichiometry. A schematic representation of the cpDNA is shown above, indicating the large and small single-copy regions (LSC and SSC, respectively) and a single copy of the inverted repeated region (IR). (**B**) Abundance of reads identified as corresponding to rearranged cpDNA molecules (see supplemental figure 7 for a scheme of the analysis pipeline). A slight increase, up to two-fold as compared to WT, was identified in the *radA* plants.

In order to dissociate mitochondrial effects of *radA* from possible effects on the chloroplast we tried to hemicomplement the *radA* mutant with a construction corresponding to RADA fused to either a specific mitochondrial targeting sequence (AOX1:RADA) or to a plastid targeting sequence (RBCS:RADA), under control of the *RADA* promoter (Figure 9A). These targeting sequences had been shown to be sufficient for specific and efficient organellar targeting (Carrie et al., 2009). Such constructions were used to transform heterozygote *radA-1* plants, and homozygote plants were selected (AOX1:RADA and RBCS:RADA, respectively, Figure 9B). None of the constructions could achieve complete complementation of the growth defect phenotype. We could not compare the transgenes expression levels, as the transgene-deriving proteins could not be detected with HA-specific antibodies on total plant extracts, from either leaves or inflorescences. Nevertheless, AOX1:RADA plants displayed less severely reduced growth than RBCS:RADA plants (Figure 9B), and not the deformed leaves phenotype characteristic of non-complemented *radA-1* homozygote mutants segregated at the T2 generation from heterozygote plants. Analysis of the accumulation of cross-over product L-1/2 also revealed decreased accumulation in AOX1:RADA as compared to *radA-1* or RBCS:RADA plants (Figure 9C). Taken together, the data from hemicomplemented plants also suggests that the severe growth phenotype of *radA* is mainly because of the effects on mitochondrial genome stability, and not because of chloroplast dysfunction and indirect effects on mitochondria. But RBCS:RADA plants also seemed slightly less affected than the non-complemented mutant (Figure 9B), implying that there is also a plastidial component in the growth defect phenotype of *radA*.

**Figure 9.**
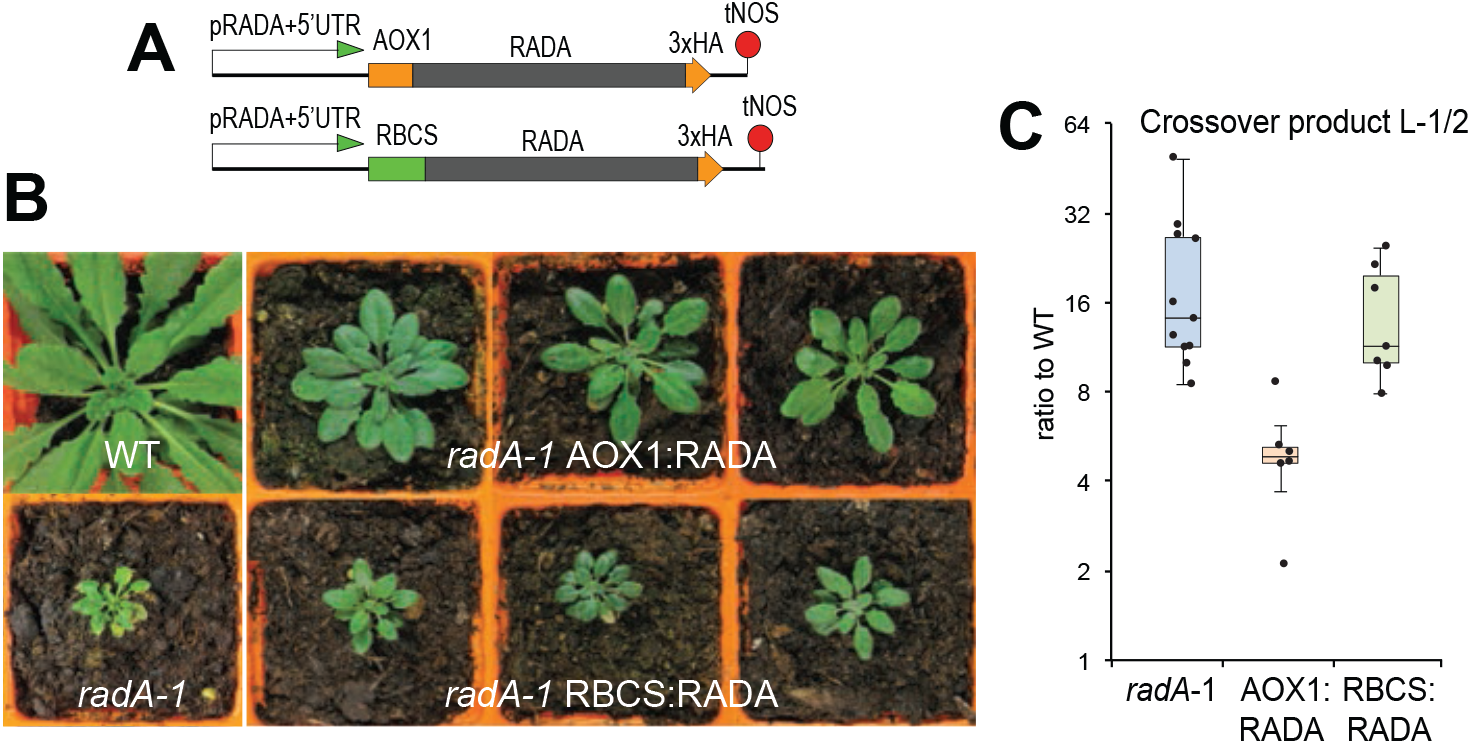
Hemicomplementation of *radA*. (**A**)Schematic representation of the constructs used for testing hemicomplementation of the *radA* phenotypes, with a construct only targeting RADA to mitochondria, by fusion to the targeting sequence of alternative oxidase (AOX1) and with a construct only targeting RADA to plastids, by fusion to the targeting sequence of the small subunit of Rubisco (RBCS). (**B**) Growth phenotypes of four-week-old hemicomplemented plants in a segregating T2 population as compared to WT and *radA-1*. Only partial complementation was observed with the AOX1:RADA construct, but plants complemented with the RBCS:RADA construct showed little or no complementation. (**C**) Relative quantification of the accumulation of crossover product L-1/2, resulting from ectopic recombination involving mitochondrial pair of repeats L in *radA-1* and in hemicomplemented plants (log2 scale).

### The *radA* mutation is synergistic with *recA3* but not with *recG1*

In bacteria the *radA* mutation is highly synergistic with *recG*. We therefore tested the epistatic relation between Arabidopsis *RADA* and *RECG1,* in *recG1 radA* double mutants. To compare all mutant plants at the first homozygous generation, heterozygous *recG1-2* plants [KO for *RECG1*, (Wallet et al., 2015)] were crossed with *radA-1* heterozygous plants used as pollen donor. In the segregating F2 generation we obtained WT, *recG1-2, radA-1* and *recG1-2 radA-1* double homozygous mutants. In this cross, plants inherit the organellar genomes of accession Ws, which is the genetic background of *recG1-2.* As described, the *recG1-2* single mutants were similar to WT (Wallet et al., 2015). The *radA-1* single mutants developed the same growth defects observed in the Col-0 background. But surprisingly the *recG1-2 radA-1* double mutants were as severely affected in growth as *radA-1* plants, with no evidence of a negative epistasis between the two mutations (Figure 10A). The double mutants were tested for their cumulative effects on mtDNA stability. In the genetic background of the Ws mtDNA the effect of the *recG1* KO mutation was mainly the accumulation of the episome resulting from the recombination across repeats EE (Wallet et al., 2015). We tested the accumulation of this episome in *recG1-2,* in *radA-1* and in the *recG1-2 radA-1* double mutant (Figure 10B). In the first homozygous mutant generation we found mild and equivalent effects of both *recG1-2* and *radA-*1 on the accumulation of the EE episome, 3 fold and 3.3 fold more abundant than the flanking mtDNA regions, respectively. However, in *recG1-2 radA-1* there was a significant increase in copy number of the EE episome, with roughly a 10 fold higher copy number than the flanking regions. Thus, at the molecular level the *recG1-2 radA-1* double mutation has an additive impact on the correct segregation of mtDNA sequences. But contrarily to what was observed in bacteria, in plants the *recG1 radA* double mutants are not more severely affected in growth than *radA* plants, implying that RADA contributes to a higher extent to mitochondrial genome maintenance than RECG1.

**Figure 10.**
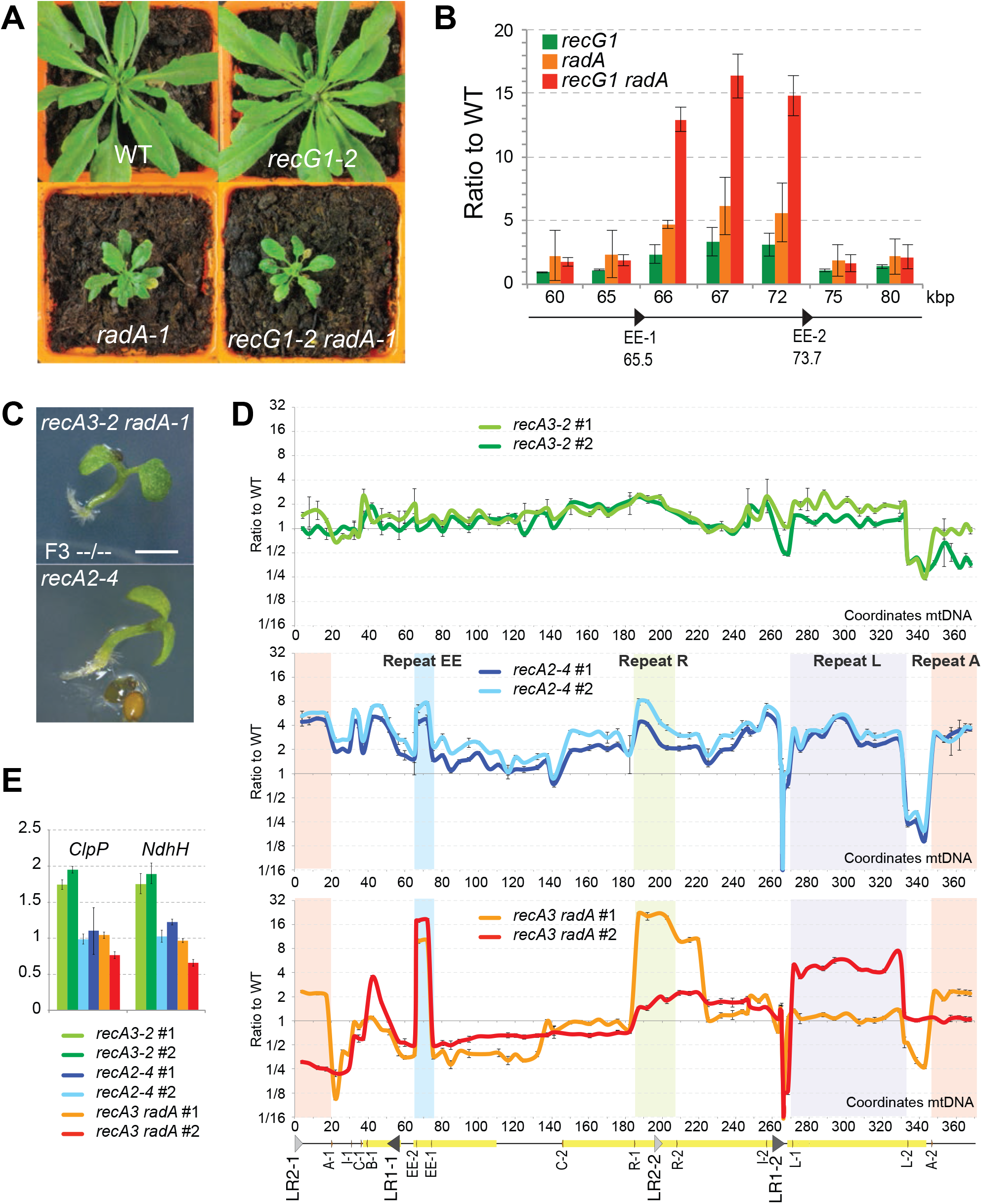
Synergistic effects of the *radA* mutation on *recG1* and *recA3*. (**A**) Crosses of *recG1-2* and *radA-1* (pollen donor) show that double homozygote *recG1 radA* plants are not more affected in their development than simple homozygote *radA* plants. (**B**) qPCR analysis of the copy number of mtDNA sequences around and within the region comprised between the pair of repeats EE, previously shown to generate an episome by recombination in *recG1* plants (Wallet et al., 2015). Autonomous replication of the episome is significantly increased in the *recG1 radA* double mutant. Results show the mean and *SD* error bars from 2 or 3 (for *recG1 radA*) biological replicates. (**C**) Double homozygote *recA3 radA* seedlings do not grow roots and do not grow further. The phenotype is similar to the one observed for *recA2* mutants (Miller-Messmer et al., 2012). The scale bar is 1 mm. (**D**) Scanning of the copy numbers of the different mtDNA regions, as described in Figure 7, in pools of *recA3 radA* double mutant seedlings. Results were compared to those of WT seedling of same size, and are represented in log scale. Sequences that are also present in the nuclear genome of Col-0 are shaded yellow in the mtDNA representation.

It was previously shown that *recG1* is synergistic with mutants deficient for *RECA3* (Wallet et al., 2015), which encodes a mitochondrial RecA ortholog that is dispensable, contrarily to RECA2 (Miller-Messmer et al., 2012). Therefore we also tested the epistatic relationship between *radA* and *recA3.* Heterozygous *recA3-2* and *radA-1* mutants (both in Col-0 background) were crossed, but no double homozygous mutants could be retrieved from F2 plants growing on soil. Seeds from sesquimutant *recA3*−/− *radA*+/− were germinated *in vitro* and seedlings were genotyped, revealing that *recA3 radA* double homozygous mutants can germinate (Figure 10C), but are unable to grow roots and to expand their cotyledons. The seedling lethal phenotype was similar to the one observed for *recA2* (Miller-Messmer et al., 2012), and also for certain later generation *radA* plants, as shown in Figure 7A. Molecular analysis showed that mtDNA stability was greatly affected. While segregating *recA3* single mutant plants showed just mild effects, an overall reduction in copy number as compared to WT seedlings of the same size was observed for the mtDNA of *recA3 radA*, some regions being more than 20 fold reduced or increased as compared to neighboring sequences (Figure 10D). The profiles were equivalent to the one obtained for *recA2* plants, both *recA2* and *recA3 radA* showing dramatic reduction of the regions between repeats L-2 and A-2 and between Large repeat 1-2 and repeat L-1. These sequences are duplicated in the nuclear genome of Col-0 (Stupar et al., 2001) and most probably the reduction corresponds to a complete loss from the mtDNA. A small sequence (278 bp) downstream LR1-2 is unique to the mtDNA, and that sequence is virtually absent in both *recA2* and *recA3 radA.* In contrast, the copy number of the cpDNA was little affected, apart for one of the *recA3 radA* replicates in which a small decrease of about 0.7-fold was observed (Figure 10E). Thus, the seedling lethality can be explained by massive problems in the replication and segregation of the mtDNA. The synergistic effect of the *radA* and *recA3* mutations suggests that the two factors intervene in alternative pathways, and that compromising both of them leads to a defect equivalent to the loss of the main RECA2 dependent recombination pathways.

### Effects of mtDNA instability on the mitochondrial transcriptome

As discussed above, the instability of the mtDNA in *radA* plants correlated with the severity of the growth defect phenotypes. But in no case we found a significant reduction in copy number of an mtDNA gene, linking the phenotype to a defect in mtDNA gene expression. In order to understand the reason for the *radA* growth defects the relative abundance of most mitochondrial gene transcripts and rRNAs was quantified by RT-qPCR. Surprisingly, no apparent defect in mtDNA gene expression was found. Rather, for most transcripts an increased accumulation was observed, as compared to WT plants of the same size (Figure 11A), up to 8-fold in the case of the *rps4* transcript. Because reshuffling of the mtDNA sequences by recombination can change the expression of non-coding ORFs and/or lead to the expression of chimeric genes we also tested the expression of a few mtDNA ORFs whose transcription could be affected by recombination across IRs. These included *orf195*, a chimera containing the sequence encoding the N-terminus of RPS3 as a result of recombination across repeats A (Figure 11B), *orf315*, which contains the sequence encoding the N-terminus of ATP9 following recombination across repeats G, as well as *orf262a* and *orf255*, which might be transcriptionally activated by recombination involving repeats E, L or H. An 8-fold increase in the abundance of *orf195* and *orf315* transcripts was observed, whose expression is driven by the *rps3* and *atp9* promoters, respectively. Thus, one of the possible reasons for the *radA* growth defects could be the expression of toxic ORFs generated by recombination that could interfere with the assembly or activity of OXPHOS complexes. However, analysis of the later on Blue-Native gels failed to detect any major problem (Figure 11C). Activity staining of Complex I and immunodectection of Complex III also revealed complexes of the same size and abundance as in WT mitochondria.

**Figure 11.**
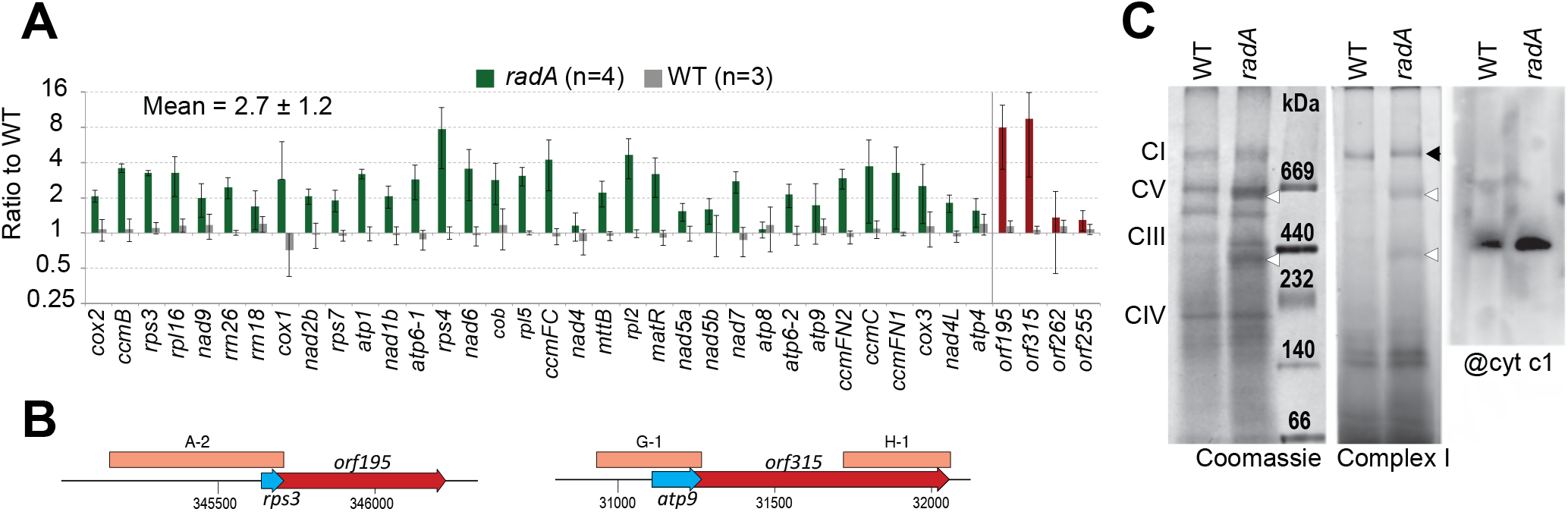
Accumulation of mitochondrial transcripts in *radA*. (**A**) Representative mitochondrial transcripts were quantified by RT-qPCR, from the RNA of 10-day-old seedlings grown *in vitro,* and normalized against a set of nuclear housekeeping genes. The quantification of several orf transcripts whose expression could be potentially affected by ectopic recombination involving IRs is shown on the right, in red. Results are on a log2 scale and are the mean from four biological replicates (two pools of *radA-1* and two pools of *radA-2* seedlings) with the corresponding *SD* error bars. (**B**) Chimeric *orf195* and *orf315* created by recombination whose transcription is augmented in *radA* plants. The regions corresponding to *rps3* and *atp9* sequences are shown in blue. Repeated sequences are represented by orange bars. Coordinates on the mtDNA are indicated. (**C**) Blue-Native gel analysis of WT and *radA* purified mitochondria. 80 μg of proteins were loaded in each well, and run in parallel with size markers (High Molecular Weight Calibration Kit, GE Healthcare). Coomassie staining, complex I activity staining and immunodetection of complex III (with a cytochrome c1-specific antibody) revealed no apparent changes in *radA* respiratory complexes. The white arrowheads indicate green-yellow bands that were already visible before staining and likely corresponded to plastidial contamination. The black arrowhead indicates the complex I activity band.

### Instability of the mtDNA in *radA* mutants affects cell cycle progression

Scanning electron microscopy (SEM) showed that *radA* epidermal leaf cells were much enlarged as compared to cells of WT leaves, suggesting an inhibition of cell division (Figure 12A). The *radA* leaves also had significantly fewer stomata (384.mm^−2^ in *radA-1 versus* 779.mm^−2^ in WT, X^2^ test=8.5E-14). To test such a possibility, the nuclear DNA ploidy levels were measured by flow cytometry, in the fully developed first true leaves of 20-day-old *radA* and WT plants (Figure 12B and C). The *radA* leaves displayed a higher proportion of 4C and 8C nuclei than WT leaves, and an important reduction in 16C nuclei, suggesting that endoreduplication is inhibited in *radA*. In all *radA* samples an accumulation of intermediate peaks (ex. 8-16C) was observed, but not in WT, also suggesting a blockage of cell cycle progression in S phase, or an induction of programmed cell death. To further explore whether mtDNA instability in *radA* impacts cell cycle progression we quantified nuclear DNA replication in root tips, using the thymidine analog 5-ethynyl-2’-deoxyuridine (EdU). In *radA* the ratio of EdU-positive nuclei was significantly reduced (Figure 12D). Counting of mitotic events in root tips also showed much reduced numbers of cells undergoing mitosis in *radA* (Figure 12E).

**Figure 12.**
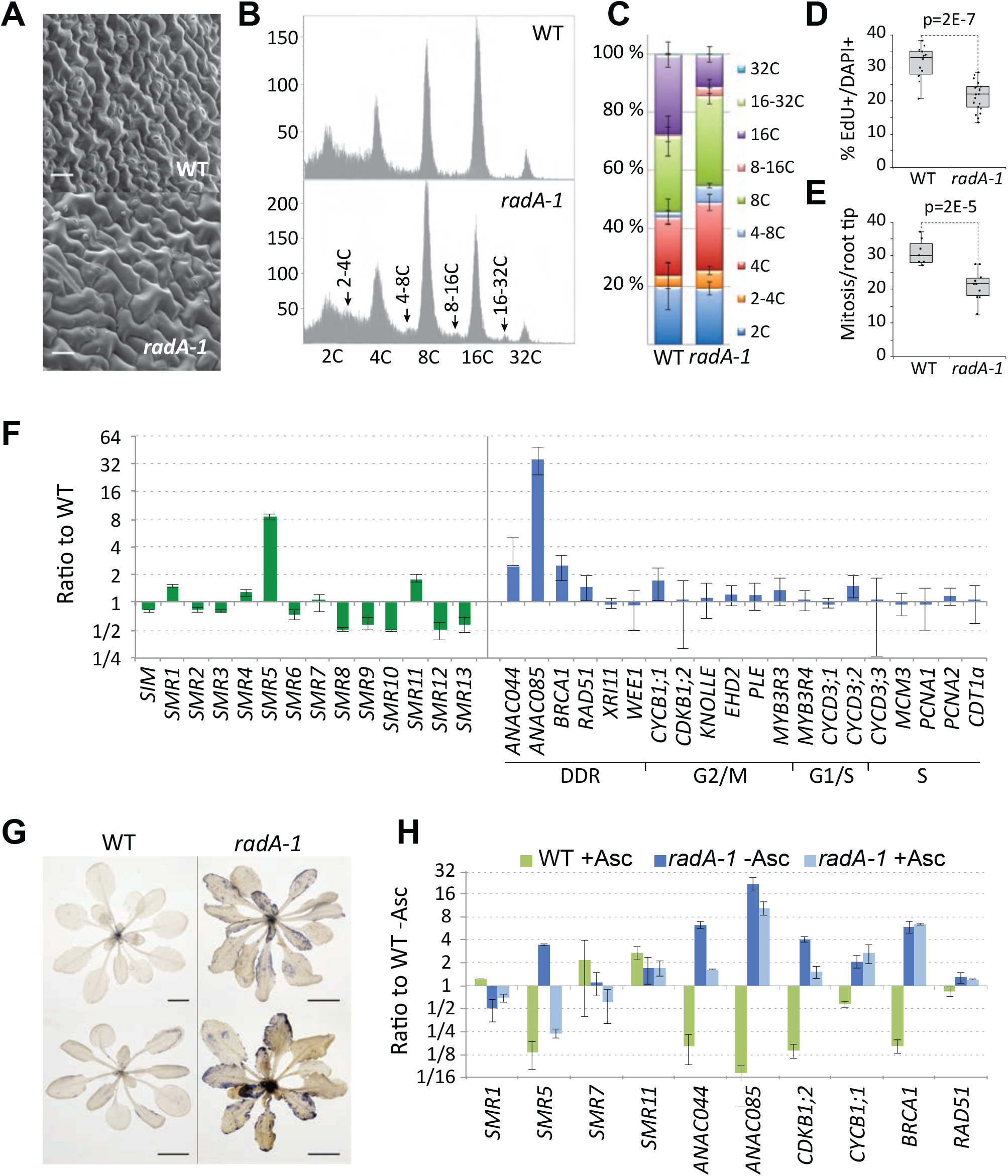
Cell cycle progression is impaired in *radA* plants. (**A**) Scanning electron microscopy images showing much larger epidermal cells and fewer stomata in *radA* leaves. Scale bar is 20 μm. (**B**) Flow-cytometry profiles obtained in WT (Col-0) and *radA.* The DNA content of nuclei extracted from the first true leaves of 20-day-old plants was analyzed. (**C**) Ploidy distribution, showing decreased endoreduplication in the mutant, with an increased proportion of 4C and 8C nuclei and a decreased proportion of 16C nuclei. Values are the average ± *SD* of 4 experiments for WT and six experiments for *radA* (n>20 000 nuclei). (**D**) Decreased DNA synthesis in the nuclei of *radA* root tip cells, as evaluated by the ratio between EdU positive and DAPI (4’,6’-diamidino-2-phénylindol) positive cells. (**E**) Decreased number of cells undergoing mitosis. Significances were calculated by Student’s t test. (**F**) RT-qPCR analysis of the expression of a set of cell cycle related genes in 10-day-old WT and *radA* seedlings, revealing significant activation of *SMR5* and *ANAC085*. The data is represented in a log2 scale and is the mean ± *SD* of three biological replicates (two pools of seedlings from *radA-1* and one from *radA-2*). (**G**) NBT (nitro blue tetrazolium) staining for O^2−^ in plants of equivalent size grown under the same conditions, showing that *radA* mutants accumulate much more ROS than WT plants. Scale bar is 1 cm. (**H**) The expression of several cell cycle-related genes is suppressed by the ROS quencher ascorbic acid (1 mM), both in WT and in *radA* 10-day-old seedlings.

**Figure 13.**
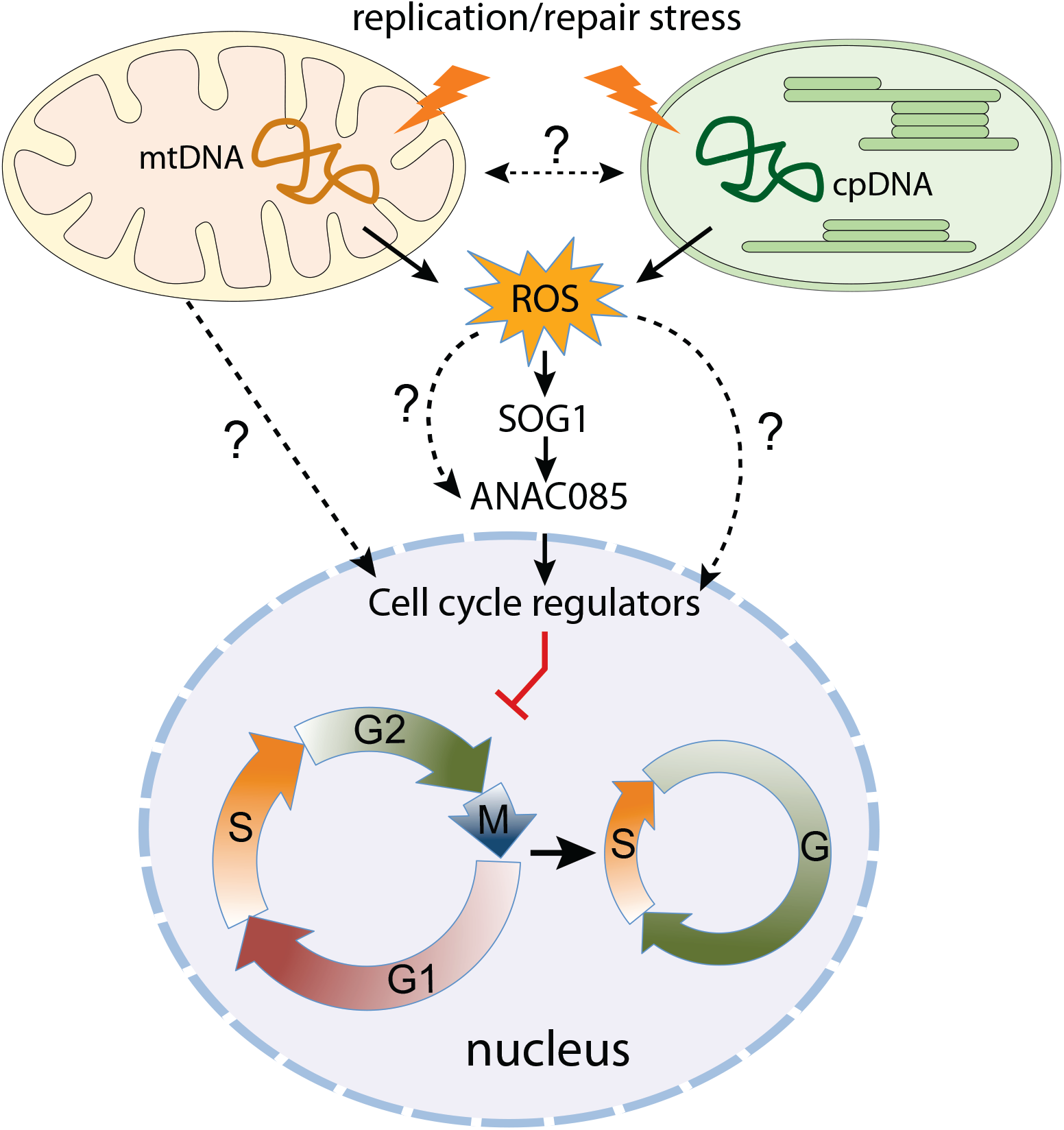
Model for possible retrograde regulation by organellar DNA instability. Possible retrograde effect of the instability of organellar genomes on the expression of cell-cycle regulators inhibiting plant growth and development. The release of ROS from mitochondria and/or chloroplast could be the signal transmitted to the nucleus.

It is known that cell cycle checkpoints adjust cellular proliferation to changing growth conditions, arresting it by inhibiting the main cell cycle controllers that include cyclin-dependent kinases (CDKs) (Nowack et al., 2012). In plants, *SMR* genes encode inhibitors of CDK-cyclin complexes that are transcriptionally induced in response to changing conditions, integrating environmental and metabolic signals with cell cycle control (Dubois et al., 2018; Hudik et al., 2014; Yi et al., 2014). Interestingly, chloroplastic defects have been shown to induce cell cycle arrest through the induction of *SMR5* and *SMR7* (Duan et al., 2020; Hudik et al., 2014). How these genes are activated remains to be fully elucidated, but the mechanism could involve the activation of the DNA damage response (DDR). This signaling cascade is controlled by the ATM and ATR kinases (Abraham, 2001) that phosphorylate the SOG1 transcription factor, leading to the up-regulation of thousands of target genes, including *SMR5* and *SMR7* (Yoshiyama, 2016; Yoshiyama et al., 2013). We have therefore tested the expression of several cell cycle markers and of several genes that are transcriptionally activated as part of the DDR. These included 14 *SMR* genes, selected cell cycle-related genes and the NAC-type transcription factors ANAC044 and ANAC085 that inhibit cell cycle progression in response to DNA damage through their activation by SOG1, but also in response to heat stress through a SOG1-independent pathway (Takahashi et al., 2019). Among the 14 SMR genes tested we found strong induction of *SMR5* in *radA* (Figure 12F, left panel). But transcription of *SMR7*, whose activation has been described as associated with that of *SMR5* in the case of chloroplastic defects (Hudik et al., 2014; Yi et al., 2014), remained unchanged. ANAC085 was strongly activated in *radA,* more than 30-fold as compared to WT, as well as ANAC044 but to a much lower degree (Figure 12F, right panel). We also observed a low-level induction of the DDR genes *BRCA1* and *RAD51*, and of the *CYCB1;1* gene encoding a cyclin associated with G2 arrest and DSB repair (Weimer et al., 2016).

SMR5 is a cyclin-dependent kinase inhibitor that is induced by different conditions leading to oxidative stress (Peres et al., 2007). ROS-dependent transcriptional activation of *SMR5* and of *SMR7* was confirmed in several ROS-inducing conditions (Yi et al., 2014). We have therefore tested whether the induction of *SMR5* in *radA* mutants could be due to the accumulation of ROS. Whole rosettes of WT and *radA* plants of same size were stained with nitro blue tetrazolium (NBT) to reveal O^2−^, and a much higher accumulation of ROS was indeed confirmed in *radA* plants (Figure 12G). To confirm the effect of ROS on the activation of cell cycle regulators, *radA* and WT plants were grown in the presence of the ROS quencher ascorbic acid, and the expression of selected genes tested. In both WT and mutant seedlings the expression of *SMR5*, *ANAC044*, *ANAC085* and *CDKB1;2* were significantly reduced, as compared to plants grown in the absence of ascorbate, up to 8-fold in the case of *SMR5* in *radA* (Figure 12H). Thus, a component of the *radA* growth phenotype is apparently a retrograde response that activates cell cycle regulators to inhibit cell proliferation. This could be because of the release of ROS as a consequence of mtDNA instability.

## DISCUSSION

The role of RadA (or Sms) in bacterial HR has been described only very recently (Cooper et Lovett, 2016; Marie et al., 2017). RadA is a hexameric helicase loaded by RecA on either side of the D-loop, to allow hybridization of the invading ssDNA with the recipient DNA (Marie et al., 2017). Our results show that plant RADA apparently has similar activity in organelles, but a more essential role in genome maintenance than its bacterial counterpart. A previous report proposed that plant RADA is a nuclear protein (Ishibashi et al., 2006), but our results just showed RADA targeted to mitochondria and chloroplasts, where it probably localizes in nucleoids. That is consistent with the presence of predicted targeting sequences in all plant sequences.

The modeled structure of plant RADA suggested that bacterial and plant proteins have similar activities. We established this conservation of activity by showing that Arabidopsis RADA complements the survival of a bacterial *radA* mutant under genotoxic conditions, as efficiently as the *E. coli* protein when brought in *trans*. Similarly, we showed that plant RADA preferentially binds ssDNA and accelerates the *in vitro* strand-exchange reaction initiated by RecA, as described for bacterial RadA (Cooper et Lovett, 2016). As reported for bacterial RadA we found that Arabidopsis RADA is not able to initiate strand invasion, and can only promote branch-migration. However, it was previously reported that the rice ortholog is able to promote D-loop formation (Ishibashi et al., 2006). In that report a different assay system was used, based on the invasion of double-stranded supercoiled plasmid by a labeled oligonucleotide, and the efficiency seemed very low. With the Arabidopsis recombinant protein and in our test system we could not reproduce such an activity.

Several differences were observed, between the activities of Arabidopsis RADA and those of bacterial RadA. RADA could bind ssDNA-containing molecules without the need for ADP or ATP. But in the presence of ADP or ATP it formed higher molecular weight complexes with ssDNA, suggesting a higher degree of protein oligomerization. The presence of ATP activates the translocation of bacterial RadA and causes the dissociation of the protein from DNA (Marie et al., 2017). Only the mutant of the ATPase domain remains associated with DNA in the presence of ATP, because it is unable to activate translocation (Marie et al., 2017). In contrast, both the WT Arabidopsis RADA and the Walker A K201A mutant could bind to ssDNA and form high molecular weight complexes in the presence of ATP or ADP. A possible explanation could be that under our test conditions the ATPase activity of RADA is not functional, preventing translocation on the DNA. But that seems unlikely given that the recombinant protein could accelerate *in vitro* strand exchange, and should therefore have translocase activity. Thus, the conditions of interaction of RadA (or RADA) with DNA remain controversial. Marie et al. (2017) observed the binding of bacterial RadA to DNA in the absence of ATP, and translocation in the presence of ATP. Cooper and Lovett (2016) observed RadA binding to DNA only in the presence of ADP, while we observed binding of plant RADA to DNA regardless of the presence of ADP or ATP. Departing also from what was described for bacterial RadA we found that plant RADA by itself can promote branch migration of recombination intermediates, in the absence of RecA, while the interaction with RecA was described as necessary for bacterial RadA activity

In bacteria, the mutation of a single branch migration factor is only slightly deleterious for cell growth and DNA repair, and that is particularly true for RadA (Beam et al., 2002; Cooper et al., 2015). In plants however, the single loss of RADA severely affects plant development and fertility. This result also contrasts with the lack of a notable developmental phenotype observed for the Arabidopsis *recG1* mutants (Wallet et al., 2015). It therefore seems that, in plants, RADA has a more important role than RECG1. Since the RuvAB migration pathway is absent in plants, it is possible that HR in plant organelles has evolved by favoring the RADA pathway. Surprisingly, the double mutant *radA recG1* is not more affected in its development than the *radA* single mutant, while in bacteria the mutation of the different branch migration pathways is particularly synergistic (Beam et al., 2002; Cooper et al., 2015). The bacterial *radA recG* double mutant is more severely affected than the *recA* single mutant, indicating that the accumulation of unprocessed branched intermediates is more detrimental to the cell than the lack of recombination. In agreement with this hypothesis, the bacterial triple mutant *recA radA recG* is less affected than the *radA recG* double mutant. In plants, the *recA2* mutant is lethal at the seedling stage, and a double mutant *recA2 recA3* could not be segregated, suggesting a more vital role of recombination in organelles than in bacteria (Miller-Messmer et al., 2012). It is therefore surprising that the mutation of all known branch migration pathways in plants is not more deleterious than the mutation of the recombinase.

It is possible that non-processing of recombination intermediates is not as detrimental in plant organelles as in bacteria. But it can also be hypothesized that a further alternative pathway for the processing of recombination intermediates exists in plant organelles. The synergy of the *radA* and *recA3* mutations recalls that observed in the double mutants of bacterial branch migration factors (Cooper et al., 2015). Thus, it might be that RECA3 can process the recombination intermediates created by the RECA2 recombinase, thanks to its branch-migration activity that is intrinsic to RecA-like proteins (Cox, 2007). Furthermore, RECA3 is characterized by the absence of an acidic C-terminal sequence that is found in all other RecA-like proteins, including RECA1 and RECA2 (Miller-Messmer et al., 2012; Shedge et al., 2007). In bacteria, the C-terminus is a site for interaction with many other proteins that regulate RecA activity and its deletion results in a conformational change in the RecA-DNA filament that enhances almost every one of the RecA functions (Cox, 2007). In eukaryotes, RAD51 paralogs can also be involved in the regulation of recombinase functions (Chun et al., 2013; Qing et al., 2011). As an example, XRCC3, which is part of the CX3 complex, is necessary for the stabilization of heteroduplexes and controls the extent of gene conversion, thus fulfilling roles reminiscent of branch migration factors (Brenneman et al., 2002). In this respect, RECA3 might have evolved to display enhanced branch migration activity and to be partially redundant to RADA. Nonetheless, unlike RADA, RECA3 apparently retained strand invasion activity, and could partially complement the bacterial *recA* mutant in the repair of UV-induced lesions (Miller-Messmer et al., 2012). RECA3 is therefore also partially redundant with RECA2. Whereas RECA2 and RADA would have specialized in homologous sequence recognition and invasion and branch migration, respectively, RECA3 would have retained both functions.

It is also conceivable that RECA3 acts in an alternative recombination pathway independent of RECA2 and RADA. In the absence of RADA, processing of the intermediates produced by the RECA2 pathway would require activation of the alternative RECA3-dependent recombination pathway. The loss of both RADA and RECA3 would overload the system with unresolved recombination intermediates, which would be lethal for the plant. The accumulation of toxic unresolved structures could be also the cause for the sever growth inhibition observed in later generations of *radA*, as described for Figure 7A. The most affected ones showed similar seedling lethality and root growth inhibition as observed for *recA2* and *recA3 radA.*

We show here that the absence of RADA results in a significant increase in mtDNA ectopic recombination. This could be because of the repair of DSBs by error-prone break-induced replication (BIR), triggered by a deficiency in HR functions required for accurate replication-coupled repair (Christensen, 2018; Gualberto et Newton, 2017). It apparently leads to the formation of sub-genomes that replicate more rapidly and without coordination with the rest of the mtDNA, as it was observed in *recG1* for the episome resulting from recombination involving repeats EE (Wallet et al., 2015). The more the structure of the mtDNA is modified, the more the development of the plant is affected. However, no region of the mtDNA containing functional genes is lost, and all mitochondrial genes that were tested are expressed. The expression of tRNAs has not been tested, but no region comprising a tRNA gene is lost in *radA* plants. Thus, we could not correlate the developmental phenotypes with the reduced expression of a transcript for an OXPHOS subunit, or for a factor required for OXPHOS subunit synthesis and complex assembly. The BN gel analysis also did not reveal any obvious defect in OXPHOS complexes. Nevertheless, the increase in ectopic recombination can lead to the creation and expression of chimeric genes that could encode toxic proteins (Hanson et Bentolila, 2004; Touzet et Meyer, 2014).

Despite dual targeting, the absence of RADA does not seem to have deleterious effects in the chloroplast. Analysis of Illumina sequences revealed only a slight increase in reads corresponding to cpDNA rearrangements, which can hardly be correlated with the severe growth defects of *radA* plants. As discussed elsewhere, that might be because of the absence of IRs in the cpDNA of Arabidopsis that can promote genome instability in conditions of replicative stress (Gualberto et Newton, 2017). Mutants of *RECG1* and *RECA2* that are also dually targeted did not reveal either any problems in cpDNA maintenance. To dissociate the effects of RADA in mitochondria and plastids we performed hemicomplementation of the Arabidopsis *radA* mutant with the AOX1:RADA or RBCS:RADA constructs, whose products were expected to be specifically targeted to either mitochondria or plastids, respectively. Full complementation of the growth defect phenotype was not achieved, but mutant plants complemented with the AOX1:RADA construct were less affected in their growth than those complemented with RBCS:RADA. Taken together, the TEM images, the mtDNA and cpDNA genomic analyses and the hemicomplementation results all agree that RADA is particularly important in the mitochondrial compartment.

The severe developmental phenotypes elicited in *radA* mutants seem to partially result from a mitochondrial retrograde signaling that promotes inhibition of cell cycle progression. The cell cycle is an energy demanding process that can be arrested by defects in respiration or photosynthesis (Riou-Khamlichi et al., 2000). Several studies have shown that chloroplastic defects lead to altered cell cycle regulation including reduced cell proliferation and premature endoreduplication (Duan et al., 2020; Hudik et al., 2014; Pedroza-Garcia et al., 2019). That seems to be also the case in *radA,* as suggested by the increased size of epidermal cells and confirmed by determination of nuclear ploidy, analysis of EdU incorporation and mitotic activity, although in this case endoreduplication was reduced in the *radA* mutant. Our results thus show that defects in the maintenance of mitochondrial genome integrity can also interfere with cell cycle progression in plants. Consistently, expression of *RADA* is higher in proliferating tissues, although it does not seem to be cell cycle regulated according to data generated with synchronized BY-2 cells (Trolet et al., 2019), suggesting that maintenance of mitochondrial genome integrity is crucial in highly dividing cells. A reduced mitotic index has also been observed in *recA3 msh1* double mutants, which develop a sever growth defect phenotype similar to *radA* (Shedge et al., 2007). And in Drosophila it has been established that mitochondrial dysfunction activates retrograde signals, including ROS, to modulate cell cycle progression (Owusu-Ansah et al., 2008)

In the case of chloroplast dysfunction, it has been suggested that accumulation of ROS is responsible for the observed cell cycle arrest. Similarly, it is possible that mitochondrial genome instability in *radA* results in sub-optimal function of the OXPHOS complexes and in ROS production, triggering the arrest of the cell cycle and the developmental phenotypes observed. But just functional deficiency of OXPHOS complexes does not *per se* activate regulators of cell cycle, since in mutants of complex I there was no evidence for the activation of genes such as SMR5 or SMR7 (Meyer et al., 2009). In several chloroplast-deficient mutants such as *crl* or the triple *reca1why1why3*, ROS accumulation is thought to activate the DNA Damage Response master regulator SOG1, leading to cell cycle arrest and activation of DNA repair (Duan et al., 2020; Hudik et al., 2014; Pedroza-Garcia et al., 2019). Indeed, DDR mutants and particularly *sog1* are hypersensitive to genotoxins targeting organelles (Pedroza-Garcia et al., 2019) and ROS accumulation leads to SOG1 activation through its phosphorylation by the ATM kinase (Yi et al., 2014). However, recent results showed that SOG1, ATM and ATR are not required for the activation of *SMR5* and *SMR7* in the *crl* mutant, in which chloroplast homeostasis is severely compromised (Li et al., 2020), suggesting that alternative pathways exist to control cell cycle progression depending on chloroplast activity. In line with this hypothesis, we found that a number of SOG1 targets are only slightly mis-regulated in *radA* mutants. In addition, treatment with the ROS scavenger ascorbate only partially rescued the activation of genes involved in cell cycle arrest in the mutant. Interestingly, we observed that *ANAC085*, a direct SOG1 target that can also be activated directly by abiotic stress such as heat (Takahashi et al., 2019) is strongly upregulated in the *radA* mutant, and that this induction is largely independent of ROS accumulation. It is thus tempting to speculate that retrograde signaling triggered by mitochondrial genome instability also relies on alternative pathways independent of canonical DDR signaling, and only partly mediated by ROS accumulation (Supplemental Figure 8). Further work will be required to fully decipher how defects in the maintenance of organellar genomes integrity lead to an inhibition of cell cycle progression.

## METHODS

### Plant Material

Arabidopsis T-DNA insertion mutant lines, all in the Col-0 background, were obtained from the Nottingham Arabidopsis Stock Centre (*radA-1*: SALK_097880, *radA-2*: WiscDsLoxHs058_03D). Plant genotypes were determined by PCR using gene and T-DNA specific primers. Seeds were stratified for 3 days at 4 °C and plants were grown on soil or on half-strength MS medium (Duchefa) supplemented with 1 % (w/v) sucrose, at 22 °C. DNA was extracted using the cetyltrimethylammonium bromide method. RNA was extracted using TRI Reagent (Molecular Research Centre, Inc.). For RT-qPCR experiments, 5 μg of RNA were depleted from contaminating DNA by treatment with RQ1 RNase-free DNase (Promega) and were reverse-transcribed with Superscript IV Reverse Transcriptase (Thermo Fisher Scientific), according to the manufacturer’s protocol using random hexamers. For mutant complementation, the WT *RADA* gene and promoter sequence was cloned in binary vector pGWB613, fused to a C-terminal 3xHA tag, and used to transform heterozygous *radA–1* plants. Expression of the transgene in the T1 transformants was monitored by western-blot with a HA-specific antibody. For hemicomplementation, i) the promoter sequence and 5’ UTR of *RADA*, ii) the N-terminal sequence (49 codons) of soybean alternative oxidase 1 (AOX1; NM_001249237) or the N-terminal sequence (80 codons) of the small subunit of ribulose bisphosphate carboxylase (RBSC; At1g67090), and iii) the RADA sequence minus its first 86 codons fused to a C-terminal HA epitope and Nos terminator, were assembled together by MultiSite Gateway cloning in vector pB7m34GW,0 (https://gatewayvectors.vib.be/), giving constructions AOX1:RADA and RBCS:RADA respectively. These constructs were used to transform heterozygous *radA-1* plants by floral dip. For promoter:GUS fusion expression analysis, the *RADA* promoter and 5’-UTR (1114 bp) was cloned in vector pGWB633 and transformed into Col-0 plants.

### Isolation of mitochondria and Blue Native PAGE

Arabidopsis mitochondria were prepared from the aerial part of three-week old plants grown on soil, by differential centrifugation and step density gradients as described (Meyer et al., 2009). Arabidopsis mitochondrial complexes were solubilized with 1.5 % dodecylmaltoside and analysed on BN-PAGE 4-13% (w/v gels, as described (Wittig et al., 2006). Blue staining and in gel NADH dehydrogenase activity were as described (Sabar et al., 2005).

### Bioinformatics analysis

Bacterial and plant sequences were identified in the databases by BLASTP and TBLASTN. Alignments were constructed with T-Coffe implemented in the Macvector package. Targeting prediction were tested using the SUBA web site (http://suba.plantenergy.uwa.edu.au) Phylogenetic trees were built with PhyML v3.1 (www.phylogeny.fr) using the neighbor-joining method implemented in the BioNJ program. Graphical representations were performed with TreeDyn (v198.3). The Arabidopsis RADA structure was modeled on the structure of RadA from *Streptococcus pneumoniae* (pdb: 5LKM), using Modeller (http://salilab.org/modeller/about_modeller.html). For Illumina sequence analysis of the cpDNA, reads were aligned against the Arabidopsis reference genomes using Burrows-Wheeler Aligner (BWA) (Li et Durbin, 2009) and filtered to keep only those mapping to the cpDNA. The coverage was extracted with Bedtool genomecov (Quinlan et Hall, 2010) and coverage positions were rounded down to the upper kb and the coverage of each 1 kb-range was normalized relative to 1,000,000 total Arabidopsis reads. For analysis of rearranged sequences, reads properly pairing to the cpDNA were filtered to only keep those showing a short-clipping sequence (threshold 20 nucleotides) and no indel (looking for the presence of a S in the CIGAR string without any I,D and H). The short-clipping sequences were then extracted with SE-MEI/extractSoftclipped (github.com/dpryan79/SE-MEI) and aligned using bowtie2 (Langmead et Salzberg, 2012) against the cpDNA. The positions of the short-clipping sequences mapping the cpDNA and of their relative read were rounded down to the upper kb to analyse the localization of the rearrangement. Those corresponding to the isomerization that results from the recombination involving the large inverted repeats were filtered out.

### Intracellular Localization

The cDNA sequence coding the Arabidopsis RADA N-terminal domain (first 174 codons) was cloned into pUCAP-GFP, derived from pCK-GFP3 (Vermel et al., 2002), and the expression cassette under control of a double 35S promoter was transferred to the binary vector pBIN+. For full protein fusion to GFP the gene sequence and 5’-UTR (from −156 to 4172 relative to the start codon) was cloned in vector pUBC-paGFP-Dest (http://n2t.net/addgene:105107). Arabidopsis Col-0 plants were transformed by the floral dip method and leaves of selected transformants were observed on a Zeiss LSM700 confocal microscope. The fluorescence of GFP and chlorophyll was observed at 505 to 540 nm and beyond 650 nm, respectively. For mitochondrial co-localization, leafs were infiltrated with a 1/1000 dilution of MitoTraker^®^ orange (Thermo Fisher Scientific) solution. Excitation was at 555 nm and observation at 560-615 nm. Nucleoid co-localization was tested by biolistic co-transfection in *Nicotiana Benthamiana* epidermal leaf cells with a PEND:dsRED construction (Melonek et al., 2012).

### In vitro strand exchange reaction

Recombination assays were performed with single-strand linear ΦX174 virion DNA and double strand circular ΦX174 RFI DNA (New England Biolabs) linearized with PstI in 20 mM Tris-acetate pH 7.4, 12.5 mM phosphocreatine, 10 U/mL creatine kinase, 3 mM ammonium glutamate, 1 mM dithiothreitol, 2 % glycerol and 11 mM magnesium acetate. In our conditions, 20.1 μM (in nucleotides) linear single strand DNA (ssDNA), 6.7 μM RecA (New England Biolabs), 2 μM RADA are incubated with buffer for 8 min at 37 °C. Then, 20.1 μM (in nucleotides) linear double-strand DNA (dsDNA) is added and the whole reaction is incubated for 5 min at 37 °C. Finally, strand exchange is initiated by adding 3 mM ATP and 3.1 μM SSB (Merck). Aliquots are stopped at indicated times by addition of 12 μM EDTA and 0.8 % SDS. Strand exchange products were analyzed on 0.8 % agarose gels run at 4 °C in Tris-acetate EDTA buffer at 50 V and visualized after migration by ethidium bromide staining. For reactions terminated in the absence of RecA, the RecA-initiated strand exchange reaction was stopped at the indicated time and DNA was deproteinized by phenol-chloroform extraction followed by ethanol precipitation. The DNA pellet was solubilized in reaction buffer, 2 μM RADA was added and the reaction was further incubated at 37 °C for the indicated time, before quenching with 12 μM EDTA and 0.8 % SDS.

### DNA Binding Assays

For electrophoretic mobility shift assays (EMSA) the purified recombinant protein (50-500 fmol according to the experience) was incubated with oligonucleotide probes (0.01 pmol-10 fmol) 5’-radiolabeled with [y-^32^P]ATP (5000 Ci/mmol; PerkinElmer Life Science). Different dsDNA structures were prepared by annealing the radiolabeled sense oligonucleotides with a twofold excess of unlabeled complementary oligonucleotide and purified on non-denaturing polyacrylamide gels. The binding reactions were performed in 20 mM Tris-HCl pH 7.5, 50 mM KCl, 5 mM MgCl_2_, 0.5 mM EDTA, 10 % glycerol, 1 mM DTT and protease inhibitors (Complete-EDTA; Roche Molecular Biochemicals), incubated at 20 °C for 20 min and run on 8 or 4.5 % polyacrylamide gels in Tris-Borate-EDTA buffer at 4 °C. After drying gels were revealed using an Amersham Typhoon phosphorimager (GE Healthcare Life Sciences). For competition assays, labeled probe and unlabeled competitor were added simultaneously to the reaction mixture.

### Recombinant proteins

Constructs pET28-RADA minus first 48 codons and pET28A-RADA[K201A] were used to express recombinant proteins in *E. coli* Rosetta 2 (DE3) pLysS (Novagen). Transformed bacteria were grown at 37 °C until OD_600nm_ = 0.6. Cultures were then chilled to 4 °C for 30 min before addition of 0.5 mM isopropyl b-D-1-thiogalactopyranoside (IPTG) and overnight incubation at 18 °C. After growth cells were pelleted, resuspended in 50 mM Tris-HCl pH 8.0, 5 % glycerol, 300 mM NaCl, 10 mM imidazole, supplemented with 1 mM PMSF and 1 X cOmplete protease inhibitors (Merck) and lysed with a French press under 1200 PSI. The crude lysate was sonicated for 3 min, clarified by 25 min centrifugation at 17700 g and filtrated trough a Filtropur S plus 0.2 μm filter (Sarstedt). The recombinant RADA and RADA[K201A] proteins were affinity purified in a precalibrated HisTrap FF Crude (GE Healthcare Life Sciences) column run at 0.5 mL/min, washed with 50 mM Tris-HCl pH 8.0, 300 mM NaCl, 5 % glycerol, 50 mM imidazole and eluted with a 50-500 mM imidazole gradient. The recombinant protein fractions was further purified by gel filtration on Superdex S200 columns and aliquots were flash frozen in liquid nitrogen and stored at −80 °C. RADA concentration was determined by spectrophotometry using an extinction coefficient of 42,440 M^−1^ cm^−1^.

### Bacterial complementation

The *E. coli* TOP10 strain was used for routine cloning, whereas the BW25113 *radA+* and JW4352 *radA785(del)::kan* were used for complementation assays. The Arabidopsis RADA cDNA (sequence coding for amino acids 137 to 627) or the *E. coli* RadA/Sms sequence were cloned between the PstI and BamHI restriction sites of the pACYCLacZ vector (Miller-Messmer et al., 2012) under the control of the *lac* promoter. Both constructs were introduced in the JW4352 strain. The pACYCLacZ empty vector was introduced in the BW25113 and JW4352 strains as control. Bacteria were grown in LB supplemented with 10 μg/mL chloramphenicol till OD_600nm_ = 0.4 before addition of 2.5 mM IPTG. At OD_600nm_ = 1.2 bacteria were diluted 10^4^ fold and grown on LB agar plates supplemented with 15 nM ciprofloxacin, 2 mM IPTG and 10 μg/mL chloramphenicol.

### qPCR Analysis

qPCR experiments were performed in a LightCycler480 (Roche) in a total volume of 6 μL containing 0.5 mM of each specific primer and 3 μl of SYBR Green I Master Mix (Roche Applied Science). The second derivative maximum method was used to determine Cp values and PCR efficiencies were determined using LinRegPCR software (http://LinRegPCR.nl). Three technical replicates were performed for each experiment. Results of qPCR and RT-qPCR analysis were standardized as previously described (Wallet et al., 2015). The recently corrected Arabidopsis Col-0 mtDNA sequence was taken as reference (Sloan et al., 2018). Quantification of mtDNA and cpDNA copy numbers used a set of primer pairs located along the organellar genomes, as described previously (Le Ret et al., 2018; Wallet et al., 2015). Results were normalized against the UBQ10 (At4G05320) and ACT1 (At2G37620) nuclear genes. The accumulation of ectopic recombination in mtDNA was quantified using primers flanking each repeats, as described (Miller-Messmer et al., 2012). The COX2 (AtMG00160) and 18S rRNA (AtMG01390) mitochondrial genes and the 16S rRNA (AtCG00920) chloroplast gene were used for normalization. For RT-qPCR experiments the GAPDH (At1G13440) and ACT2 (At3G18780) transcripts were used as standards.

### Sequencing

Total leaf DNA of WT and *radA-1* plants was quantified with a QuBit Fluorometer (Life Technologies) and libraries were prepared with the Nextera Flex Library kit, according to manufacturer’s recommendations (Illumina) using 100 ng of each DNA sample. Final libraries were quantifieded, checked on a Bioanalyzer 2100 (Agilent) and sequenced on a Illumina Miseq system (2 × 150 paired end reads).

### Flow cytometry and EdU staining

Nuclear DNA content was measured in leaves of 20-d-old seedlings, using the CyStain UV Precise P Kit (Partec) according to the manufacturer’s instructions. Nuclei were released in nuclei extraction buffer (Partec) by chopping with a razor blade, stained with 4’,6-diamidino-2-phenylindole (DAPI) buffer and filtered through a 30 μM Celltrics mesh (Partec). Between 20,000 and 30,000 isolated nuclei were used for each ploidy level measurement using the Attune Cytometer and the Attune Cytometer software (Life Technologies). At least four independent biological replicates were analyzed. EdU staining was as described (Pedroza-Garcia et al., 2016). For each root tip (n>10), the number of mitotic events was counted directly under the microscope.

### Accession Numbers

Sequence data from this article can be found in the Arabidopsis Genome Initiative or GenBank/EMBL databases under the following accession numbers: *RADA*, At5g50340; *RECG1*, At2g01440; *RECA2*, At2g19490; *RECA3*, At3g10140; *ANAC044*, At3g01600; *ANAC085*, At5g14490; *KNOLLE*, At1g08560; *CDKB1;2*, At2g38620; *EHD2*, At4g05520; *PLE*, At5g51600; *MYB3R3*, At3g09370; *MYB3R4*, At5g11510; *CYCD3;1*, At4g34160; *CYCD3;2*, At5g67260; *CYCD3;3*, At3g50070; *MCM*2, At1g44900; *MCM3*, At5g46280; *PCNA1*, At1g07370; *PCNA2*, At2g29270; *CDT1a*, At2g31270; *CycB1;1*, At4g37490; *WEE1*, At1g02970; *BRCA1*, At4g21070; *RAD51*, At5g20850; *XRI1*, At5g48720; *SIM*, At5g04470; *SMR1*, At3g10525; *SMR2*, At1g08180; *SMR3*, At5g02420; *SMR4*, At5g02220; *SMR5*, At1g07500; *SMR6*, At5g40460; *SMR7*, At3g27630; *SMR8*, At1g10690; *SMR9*, At1g51355; *SMR10*, At2g28870; *SMR11*, At2g28330; *SMR12*, At2g37610; *SMR13*, At5g59360; AOX1, NM_001249237; RBCS, At1g67090.

## SUPPLEMENTAL DATA FILES

**Supplemental Figure 1.** Phylogenetic distribution of RADA.

**Supplemental Figure 2.** Sequences alignment.

**Supplemental Figure 3.** Tissue-specific expression of Arabidopsis RADA.

**Supplemental Figure 4.** Effect of the point mutation K201A.

**Supplemental Figure 5.** Expression, purification and characterization of recombinant RADA.

**Supplemental Figure 6.** Transmission electron microscope (TEM) images.

**Supplemental Figure 7.** Pipeline cpDNA NGS analysis.

**Supplemental Table 1.** Oligonucleotides.

## Author Contributions

NC, FW-L, CN, CR, MLR, ME and JMG performed research. NC, CR, MB, AD and JMG designed the research and analyzed data. AF analyzed the Illumina sequences. NC and JMG wrote the paper.

## Acknowledgments

We are grateful to Dr. Sandra Noir for help with flow cytometry, to Dr. Esther Lechner for vectors and technical advice and to Dr. Patrice Polard and Dr. Bénédicte Michel for helpful discussions. This work has been published under the framework of the LABEX [ANR-11-LABX-0057_MITOCROSS] and benefits from a funding from the state managed by the French National Research Agency as part of the "Investments for the future" program.

---

**Supplemental Figure 1.** Phylogenetic distribution of RADA.

Genes coding for RadA-like proteins are found in all bacteria, in land plants, green, brown and red algae, diatoms and other organisms of the Stramenopile group. *Arabidopsis thaliana,* NP_199845; *Populus trichocarpa,* EEE84551; *Vitis vinifera,* XP_002277638; *Oryza sativa,* NP_001056828; *Zea mays,* NP_001170708; *Selaginella moellendorffii,* XP_002976563; *Physcomitrella patens,* XP_001757578; *Ostreococcus tauri,* XP_003084463; *Coccomyxa subellipsoidea,* XP_005643141; *Chlorella variabilis,* EFN56488; *Cyanidioschyzon merolae,* XP_005536634; *Galdieria sulphuraria,* XP_005709405; *Ectocarpus siliculosus,* CBJ25917; *Phaeodactylum tricornutum,* XP_002178713; *Thalassiosira oceanica,* EJK58798; *Saprolegnia diclina,* EQC30001; *Phytophthora infestans,* XP_002904225; *Myxococcus xanthus,* YP_629513; *Rickettsia prowazekii,* WP_014607237; *Neisseria meningitidis,* WP_002258526; *Escherichia coli,* WP_001458566; *Microcystis aeruginosa,* WP_002742386; *Synechocystis sp,* WP_009633429; *Geminocystis herdmanii,* WP_017296205; *Bacillus anthracis,* NP_842650; *Amphibacillus jilinensis,* WP_017473696.

**Supplemental Figure 2.** Sequence alignment.

Sequence alignment between representative land plant RADA sequences and RadA from proteobacteria and cyanobacteria. The Zinc-finger and KNRFG RadA-specific motif are shaded in yellow and blue respectively, and the Walker A and B motifs in green.

**Supplemental Figure 3.** Tissue-specific expression of Arabidopsis *RADA*.

(**A**) Results extracted from Genevestigator (https://genevestigator.com/) (**B**) promoter-GUS fusion results, showing predominant expression in very young leaves, in sepals, in anthers filament and in the stigmata.

**Supplemental Figure 4.** Effect of the point mutation K201A.

Effect of the point mutation K201A (amino acids in green) on the Walker A domain of RadA in the binding and hydrolysis of ATP. The structure shown is the one from bacterial RadA (Marie et al. 2017), with bound ADP and Mg^2+^ ion (sphere). The two phosphate groups of ADP are in red.

**Supplemental Figure 5.** Expression, purification and characterization of recombinant RADA.

The Arabidopsis RADA sequence minus the first 48 codons corresponding to the OTS was cloned in the expression vector pET28a fused to a N-terminal His-tag. The recombinant RADA and Walker mutant K201A were expressed in the Rosetta(DE3) strain and purified by affinity and gel filtration. (**A**) Coomassie gel staining analysis of the recombinant proteins. (**B**) Gel filtration on Superdex S200 showed that RADA purified as two peaks of high molecular weight. (**C**) Dynamic light scattering of the protein fraction from peak 2 shows that it is monodispersed and corresponding to a size of about 340 kDa, which is consistent with a hexameric RADA molecule. (**D**) EMSA analysis of the binding to an ssDNA oligonucleotide. Fractions corresponding to both peaks give complexes of the same size, although fractions of peak 1 give predominantly higher molecular weight complexes.

**Supplemental Figure 6.** Transmission electron microscope (TEM) images

TEM images of cells from leaves of same size showed morphologically normal chloroplasts (cp) in *radA.* Mitochondria (mt) were enlarged and less electron dense as compared to mitochondria from WT cells.

**Supplemental Figure 7.** Pipeline cpDNA NGS analysis.

Schematic representation of the analysis workflow of Illumina sequences for identification of cpDNA rearrangements. Blue background boxes represent text manipulation steps, while yellow and red background ones symbolize quality filtering and mapping, respectively. Sequencing was performed on both ends of DNA fragments (R1 and R2). BWA: Burrows-Wheeler Aligner. R1 & R2: Paired-end sequencing read1 and read2.

