## Supplemental Figures for "RADA is a main branch migration factor in plant mitochondrial recombination and its defect leads to mtDNA instability and cell cycle arrest"

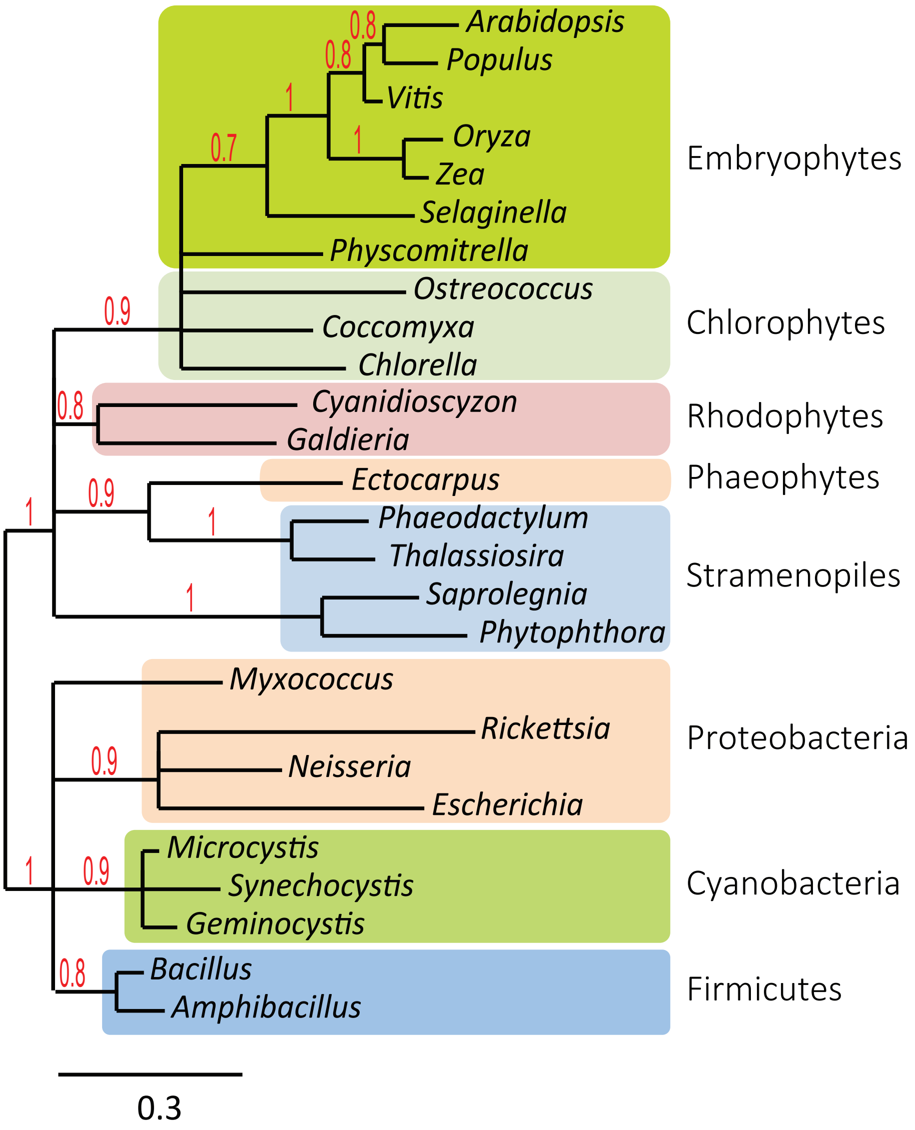


**Supplemental Figure 1.** Phylogenetic distribution of RADA.

Genes coding for RadA-like proteins are found in all bacteria, in land plants, green, brown and red algae, diatoms and other organisms of the Stramenopile group. *Arabidopsis thaliana,* NP_199845; *Populus trichocarpa,* EEE84551; *Vitis vinifera,* XP_002277638; *Oryza sativa,* NP_001056828; *Zea mays,* NP_001170708; *Selaginella moellendorffii,* XP_002976563; *Physcomitrella patens,* XP_001757578; *Ostreococcus tauri,* XP_003084463; *Coccomyxa subellipsoidea,* XP_005643141; *Chlorella variabilis,* EFN56488; *Cyanidioschyzon merolae,* XP_005536634; *Galdieria sulphuraria,* XP_005709405; *Ectocarpus siliculosus,* CBJ25917; *Phaeodactylum tricornutum,* XP_002178713; *Thalassiosira oceanica,* EJK58798; *Saprolegnia diclina,* EQC30001; *Phytophthora infestans,* XP_002904225; *Myxococcus xanthus,* YP_629513; *Rickettsia prowazekii,* WP_014607237; *Neisseria meningitidis,* WP_002258526; *Escherichia coli,* WP_001458566; *Microcystis aeruginosa,* WP_002742386; *Synechocystis sp,* WP_009633429; *Geminocystis herdmanii,* WP_017296205; *Bacillus anthracis,* NP_842650; *Amphibacillus jilinensis,* WP_017473696.

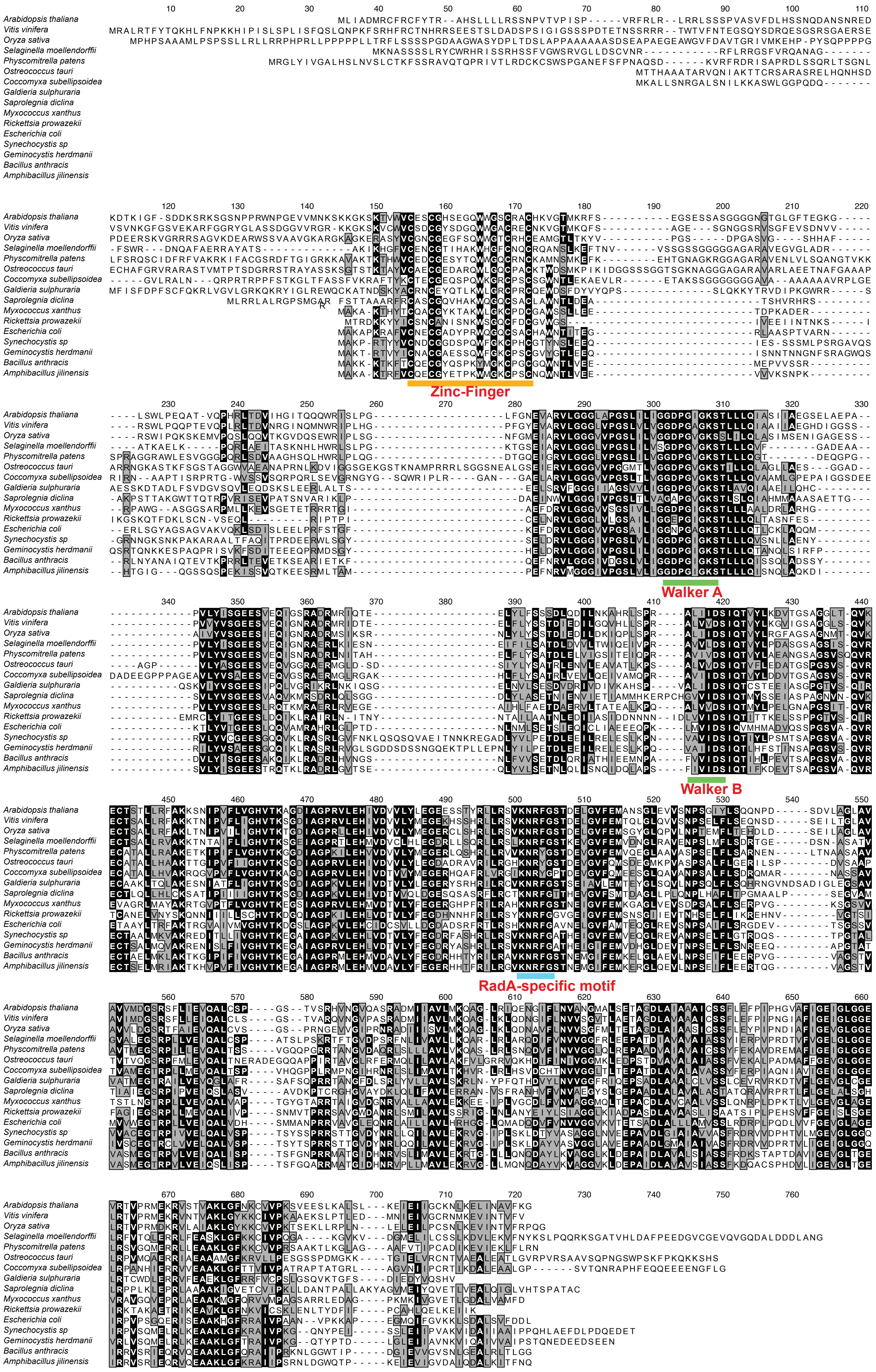


**Supplemental Figure 2.** Sequence alignment.

Sequence alignment between representative land plant RADA sequences and RadA from proteobacteria and cyanobacteria. The Zinc-finger and KNRFG RadA-specific motif are shaded in yellow and blue respectively, and the Walker A and B motifs in green.

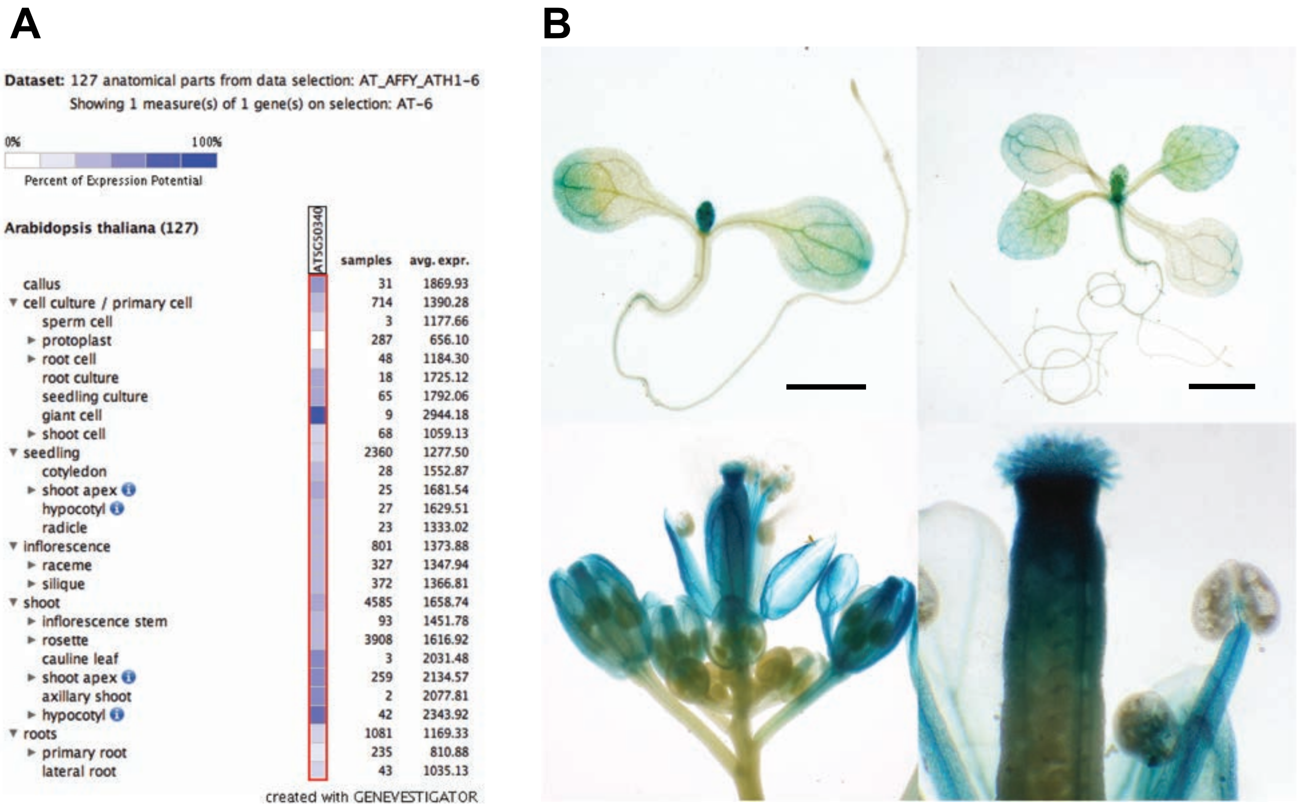


**Supplemental Figure 3.** Tissue-specific expression of Arabidopsis *RADA*.
(**A**) Results extracted from Genevestigator (https://genevestigator.com/) (**B**) promoter-GUS fusion results, showing predominant expression in very young leafs, in sepals, in anthers filament and in the stigmat. The scale bar is 1 mm.

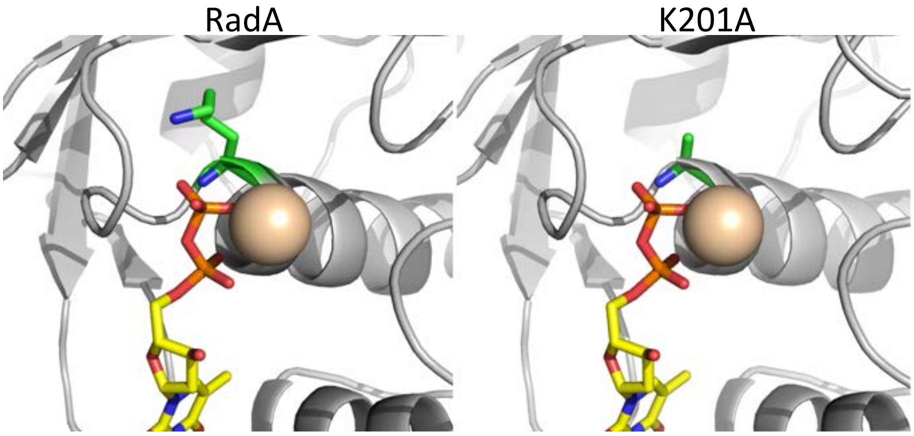


**Supplemental Figure 4.** Effect of the point mutation K201A.

Effect of the point mutation K201A (amino acids in green) on the Walker A domain of RadA in the binding and hydrolysis of ATP. The structure shown is the one from bacterial RadA (Marie et al. 2017), with bound ADP and $Mg^{2+}$ ion (sphere). The two phosphate groups of ADP are in red.

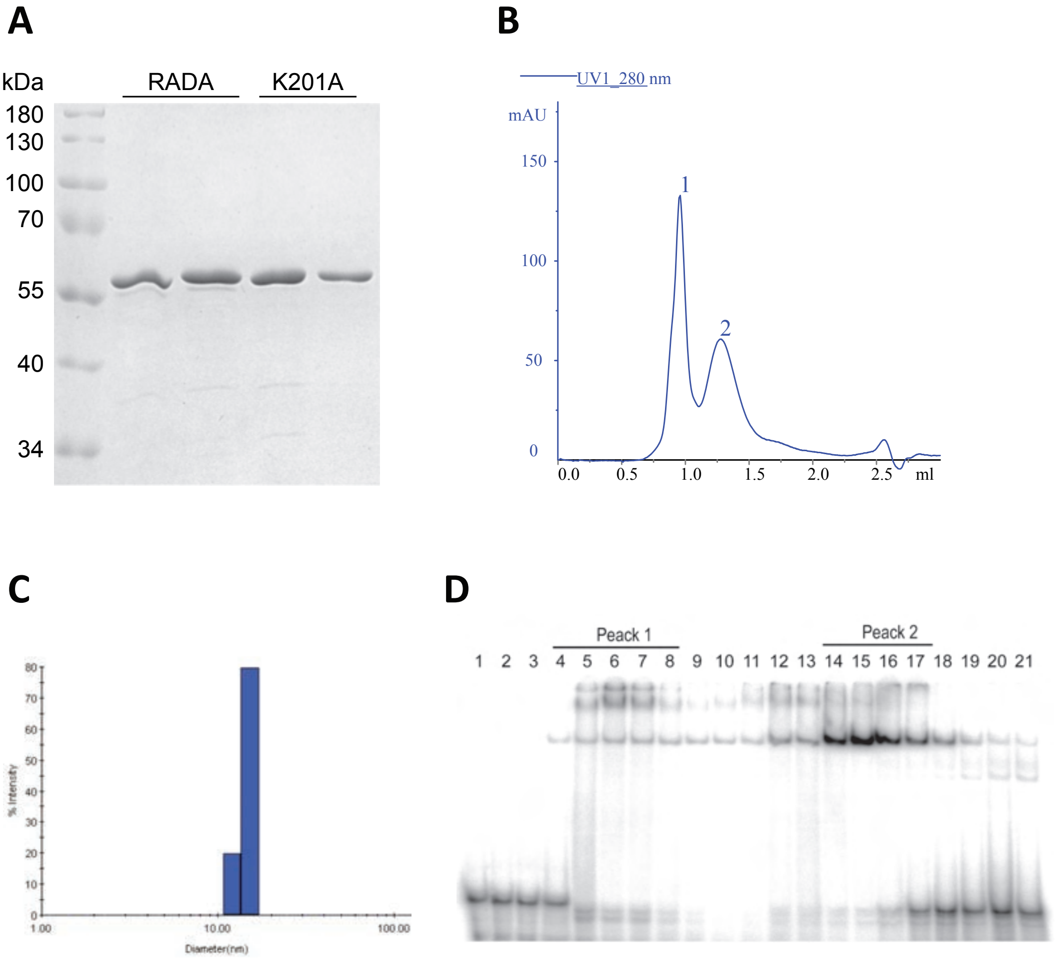


**Supplemental Figure 5.** Expression, purification and characterization of recombinant RADA. The Arabidopsis RADA sequence minus the first 48 codons corresponding to the OTS was cloned in the expression vector pET28a fused to a N-terminal His-tag. The recombinant RADA and Walker mutant K201A were expressed in the Rosetta(DE3) strain and purified by affinity and gel filtration. (**A**) Coomassie gel staining analysis of the recombinant proteins. (**B**) Gel filtration on Superdex S200 showed that RADA purified as two peaks of high molecular weight. (**C**) Dynamic light scattering of the protein fraction from peak 2 shows that it is monodispersed and corresponding to a size of about 340 kDa, which is consistent with a hexameric RADA molecule. (**D**) EMSA analysis of the binding to an ssDNA oligonucleotide. Fractions corresponding to both peaks give complexes of the same size, although fractions of peak 1 give predominantly higher molecular weight complexes.

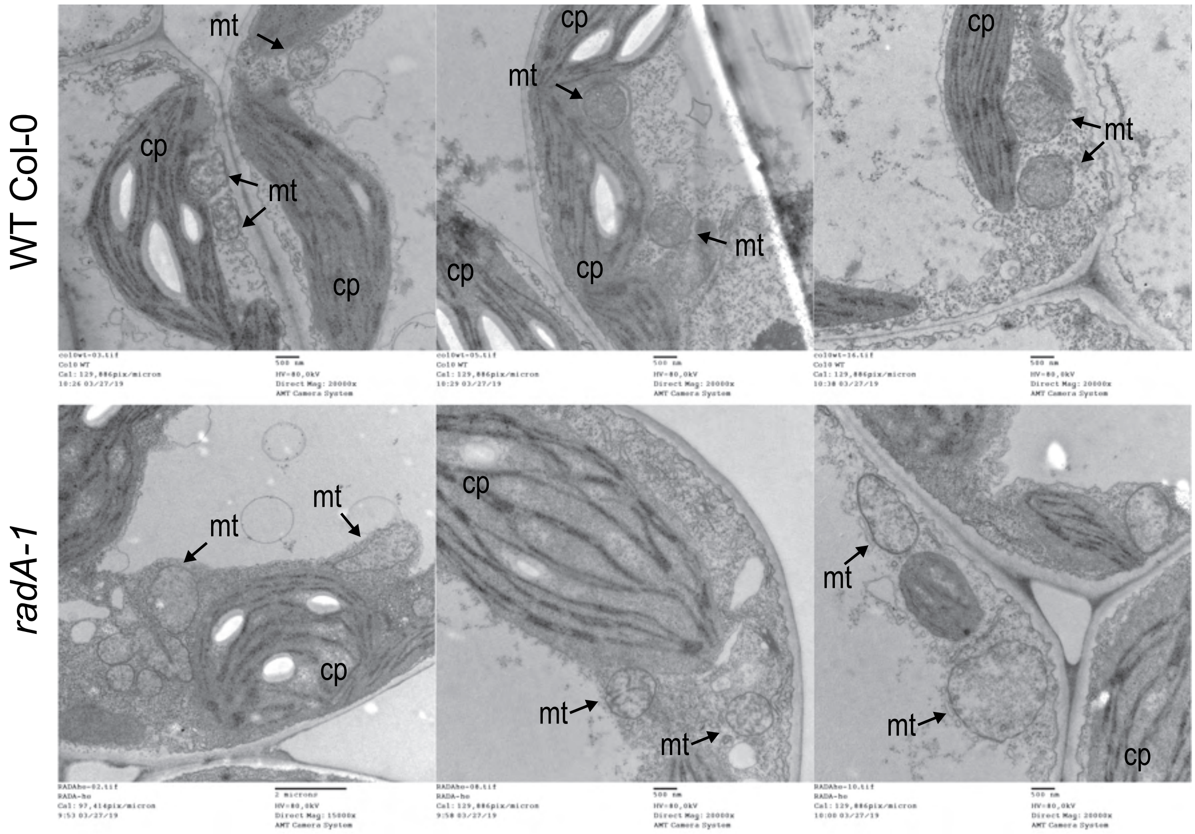


**Supplemental Figure 6.** Transmission electron microscope (TEM) images
TEM images of cells from leaves of same size showed morphologically normal chloroplasts (cp) in *radA.* Mitochondria (mt) were enlarged and less electron dense as compared to mitochondria from WT cells.

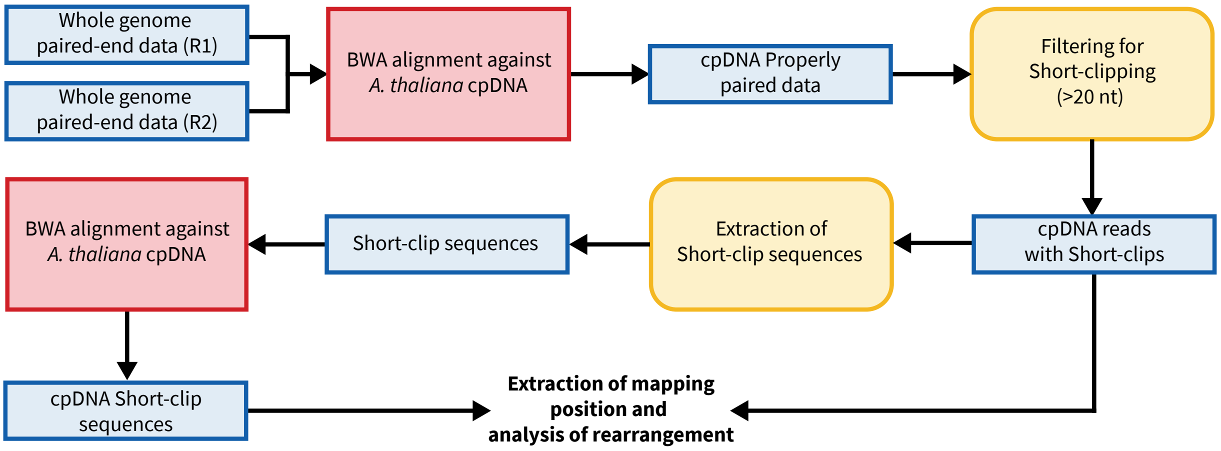


**Supplemental Figure 7.** Pipeline cpDNA NGS analysis.
Schematic representation of the analysis workflow of Illumina sequences for identification of cpDNA rearrangements. Blue background boxes represent text manipulation steps, while yellow and red background ones symbolize quality filtering and mapping, respectively. Sequencing was performed on both ends of DNA fragments (R1 and R2). BWA: Burrows-Wheeler Aligner. R1 & R2: Paired-end sequencing read1 and read2.
